# A hands-on tutorial on network and topological neuroscience

**DOI:** 10.1101/2021.02.15.431255

**Authors:** Eduarda Gervini Zampieri Centeno, Giulia Moreni, Chris Vriend, Linda Douw, Fernando Antônio Nóbrega Santos

**Author notes:** Authors Contributions: EGZC contributions encompass **Conceptualisation**, **Software**, **Writing – Original Draft Preparation** GM contributions encompass **Software**, **Writing – Original Draft Preparation** CV contribution encompasses **Writing – Review & Editing** LD contributions encompass **Conceptualisation, Writing – Original Draft Preparation**, **Supervision**FANS contributions encompass **Conceptualisation**, **Software**, **Writing – Original Draft Preparation**, **Supervision**.

## Abstract

The brain is an extraordinarily complex system that facilitates the efficient integration of information from different regions to execute its functions. With the recent advances in technology, researchers can now collect enormous amounts of data from the brain using neuroimaging at different scales and from numerous modalities. With that comes the need for sophisticated tools for analysis. The field of network neuroscience has been trying to tackle these challenges, and graph theory has been one of its essential branches through the investigation of brain networks. Recently, topological data analysis has gained more attention as an alternative framework by providing a set of metrics that go beyond pair-wise connections and offer improved robustness against noise. In this hands-on tutorial, our goal is to provide the computational tools to explore neuroimaging data using these frameworks and to facilitate their accessibility, data visualisation, and comprehension for newcomers to the field. We will start by giving a concise (and by no means complete) overview of the field to introduce the two frameworks, and then explain how to compute both well-established and newer metrics on resting-state functional magnetic resonance imaging. We use an open-source language (Python) and provide an accompanying publicly available Jupyter Notebook that uses data from the 1000 Functional Connectomes Project. Moreover, we would like to highlight one part of our notebook that is solely dedicated to realistic visualisation of high order interactions in brain networks. This pipeline provides three-dimensional (3-D) plots of pair-wise and higher-order interactions projected in a brain atlas, a new feature tailor-made for network neuroscience.

## Introduction

Neuroscience is still a young research field, with its emergence as a formal discipline happening only around 70 years ago [1]. The field has since mushroomed, and much of our current knowledge about the human brain’s neurobiology was made possible by the rapid advances in technologies to investigate the brain in vivo at high-resolution and at different scales. An example is magnetic resonance imaging (MRI), which allows us not only to measure regional characteristics of the brain’s structure non-invasively but may also be used to assess anatomical and functional interactions between brain regions [2, 3]. This expansion in the field led to an exponential increase in data size and complexity. To analyse and interpret this ‘big data’, researchers had to develop robust theoretical frameworks. Complex network science was brought to neuroscience and has been increasingly used to study the brain’s intricate communication and wiring [4, 5]. The resulting field - network neuroscience – aims to see the brain through an integrative lens by mapping and modelling its elements and interactions [4, 6].

One of the main theoretical frameworks from complex network science used to model, estimate, and simulate brain networks is graph theory [7, 8]. A graph is comprised of a set of interconnected elements, also known as *vertices* and *edges*. Vertices (also known as nodes) in a network can, for example, be brain areas, while edges (also known as links) are a representation of the connectivity between pairs of vertices [5]. Various imaging modalities are available for the reconstruction of the brain network [8, 9]. The focus of this hands-on paper will be resting-state functional MRI (rsfMRI). As the name suggests, rsfMRI indirectly measures brain activity while a subject is at rest (*i.e.*, does not perform any task). This type of data provides information about spontaneous brain functional connectivity [10]. Connectivity is often operationalised by a statistical dependency (usually a Pearson correlation coefficient) between signals measured from anatomically separated brain areas [2, 11].

Several descriptive graph metrics can then be calculated to describe the brain network’s characteristic; examples include the degree or the total number of connections of a vertex and the path length (number of intermediate edges) between two vertices [6, 12]. These metrics have consistently allowed researchers to identify non-random features of brain networks. A key example is the ground-breaking discovery that the brain (like most other real-world networks) follows a ‘small-world network’ architecture [4, 13, 14]. This refers to the phenomenon that, to minimise wiring cost while simultaneously maintaining optimal efficiency and robustness against perturbation, the brain network obeys a balance between the ability to perform local processing (*i.e.,* segregation) and combining of information streams on a global level (*i.e.,* integration).

Network neuroscience has thereby offered a comprehensive set of analytical tools to study not only the local properties of brain areas but also their significance for the entire brain network functioning. Using graph theory, many insights have been gathered on the healthy and diseased brain neurobiology [5, 9, 12, 15].

Another perspective on the characteristics of the brain network can be provided by (algebraic) topological data analysis (TDA), by analysing the interactions between a set of vertices beyond the ‘simple’ pair-wise connections (*i.e.*, higher-order interactions). With TDA, one can identify a network’s ‘shape’ and its invariant properties (*i.e.*, coordinate and deformation invariances [16, 17]). Thus, as we will illustrate along with the manuscript, TDA often provides more robustness against noise than graph theoretical analysis [18, 19], which can be a significant issue in imaging data [20–22]. Although TDA has only recently been adopted to network neuroscience, it has already shown exciting results on rsfMRI [23, 24]. For example, group-level differences in network topology have been identified between healthy subjects that ingested psilocybin (psychedelic substance) and the placebo group [25], and between attention-deficit/hyperactivity disorder children and typically developing controls [26]. A downside of this framework is that the complexity and level of abstraction necessary to apply TDA and interpret the results might keep neuroscientists without prior theoretical training from using it. Moreover, the high-order interaction structure that emerges from TDA analysis is often difficult to visualise in a realistic and understandable manner.

Therefore, we would like to facilitate the use of network neuroscience and its constituents graph theory and TDA in the general neuroscientific community by providing a step-by-step tutorial on how to compute different metrics commonly used to study brain networks as well as realistic high-order network visualisation tools. We provide a theoretical and experimental background of these metrics and include code blocks in each section to explain how to compute the different metrics. We also list several additional resources (Table 1 and Table 2) of personal preference (and by no means complete), including a Jupyter Notebook that we created to accompany this hands-on tutorial publicly available on GitHub and Zenodo [27] (see Table 1, under the Jupyter Notebooks section - Notebook for network and topological analysis in neuroscience).

**Table 1.**
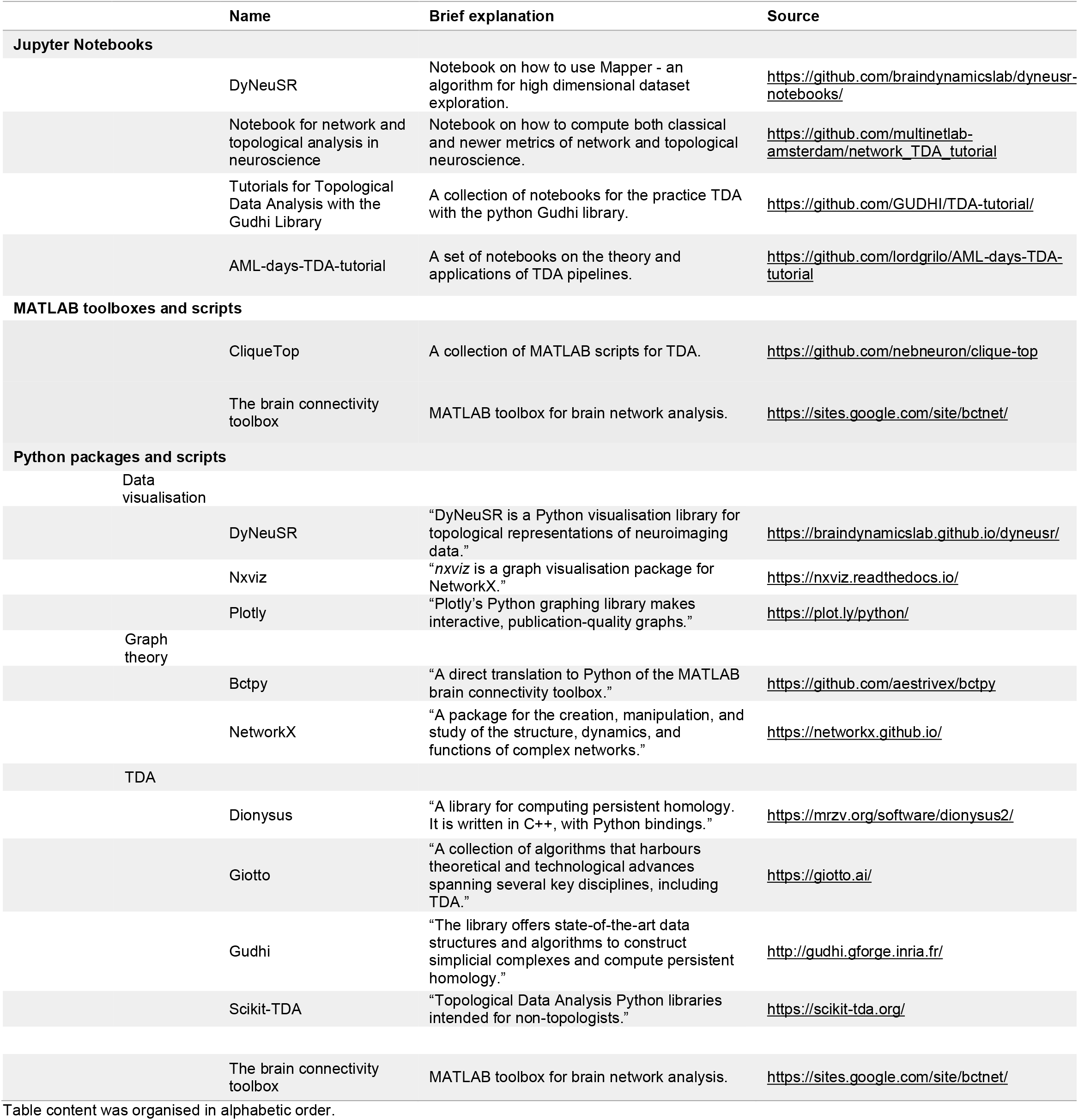
List of computational resources.

**Table 2.** List of reading resources.

| Name | Source |
| --- | --- |
| Key articles and books |  |
| Cliques and cavities in the human connectome | Reference [3] |
| Network neuroscience | Reference [4] |
| Fundamentals of brain network analysis (*the primary reference of our hands-on tutorial*) | Reference [6] |
| Graph theory approaches to functional network organisation in brain disorders: a critique for a brave new small-world | Reference [12] |
| Topology for computing | Reference [16] |
| The importance of the whole: Topological data analysis for the network neuroscientist | Reference [20] |
| Editorial: Topological Neuroscience | Reference [23] |
| What can topology tell us about the neural code? | Reference [24] |
| Homological scaffolds of brain functional networks | Reference [25] |
| Two's company, three (or more) is a simplex | Reference [28] |
| A roadmap for the computation of persistent homology | Reference [29] |
| Networks beyond pair-wise interactions: Structure and dynamics | Reference [30] |
| Clique topology reveals intrinsic geometric structure in neural correlations | Reference [31] |

Table content was organised in numerical order.

Our work differs from previous literature [12, 29] since we not only describe the concepts central to graph theory and TDA but also provide an easy-to-grasp step-by-step tutorial on how to compute these metrics using an easily accessible, open-source computer language. Furthermore, we offer new 3-D visualisations of simplicial complexes and TDA metrics in the brain that may facilitate the application and interpretation of these tools. Finally, we would like to stress that even though this tutorial focuses on rsfMRI, the main concepts and tools discussed in this paper can be extrapolated to other imaging modalities, biological or complex networks.

Since graph theory has been extensively translated for neuroscientists elsewhere, we refer the reader to the book in [6]. In this tutorial, we mainly focused on the topics covered in chapters 3, 4, 5, and the particular sections of chapters 6, 7, 8 and 9 about assortativity, shortest paths and the characteristic path length, the clustering coefficient, and modularity. In the second part of the tutorial, we explore hands-on TDA metrics, providing a summary of both theoretical and neuroscientific aspects with the calculations used in our work. We believe that our tutorial, which is far from being exhaustive, can contribute to make this emerging branch of network and topological neuroscience accessible to the reader. The codes we provide usually only requires the knowlege of the connectivity matrix and, for our realistic 3d visualisation of simplicial complexes, we only need the coordinates of the nodes of a given brain atlas. Therefore, we believe that our scripts can be adapted to different databases, image modalities and brain atlas. A short glossary with the key terms to understand this manuscript can be found in Table 3.

**Table 3.**
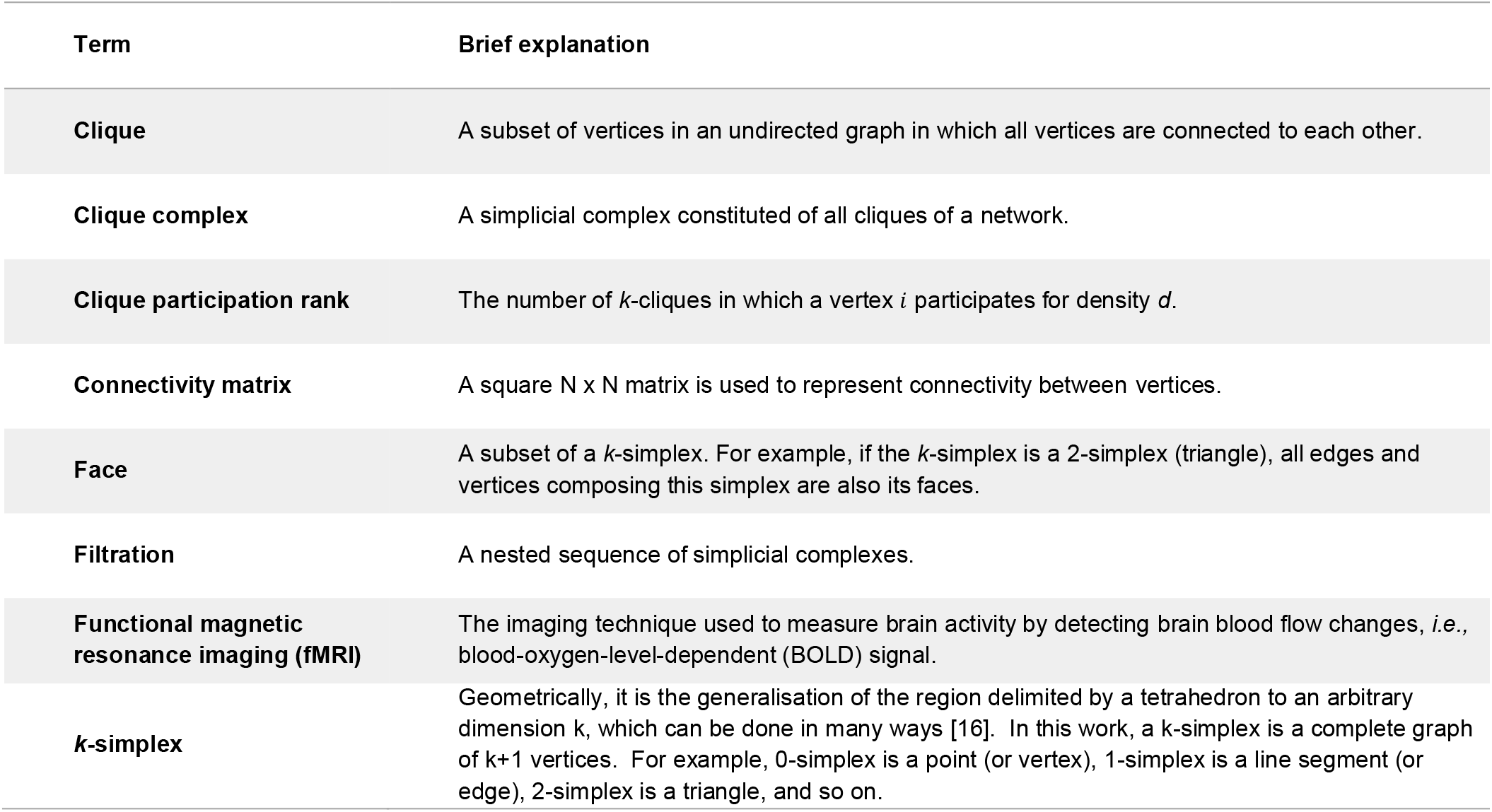
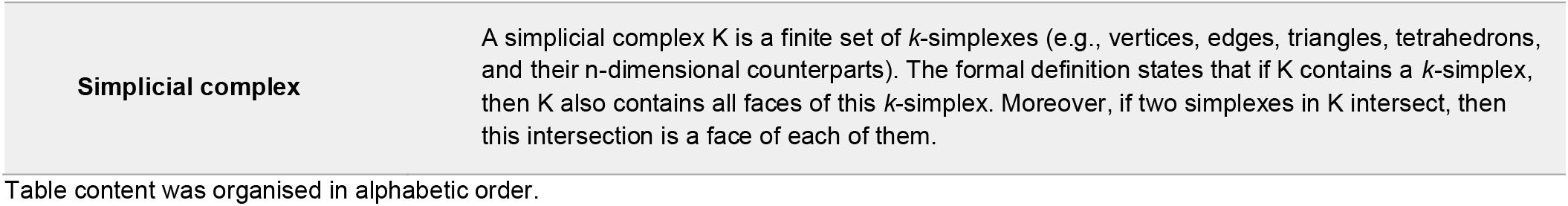
Glossary with key terms.

## Hands-on tutorial

### General requirements

The following Python 3 packages are necessary to perform the computations presented below. The accompanying Jupyter Notebook can be found on GitHub (Table 1) or Zenodo [27].

#### Code example

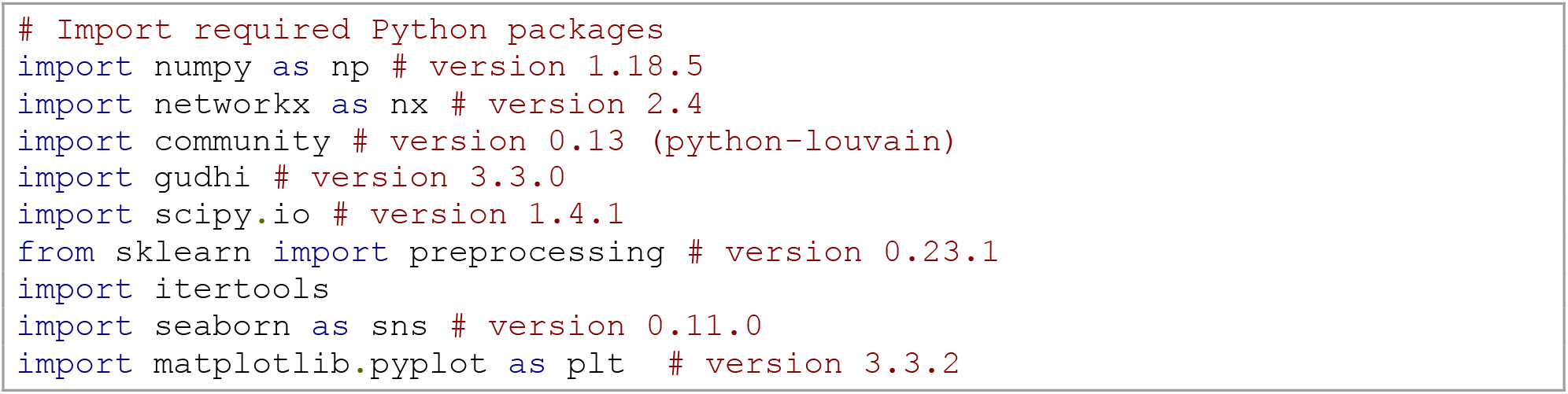

### The basis: the adjacency matrix

The basic unit on which graph theory and TDA are applied in the context of rsfMRI in our work is the adjacency or connectivity matrix (Fig 1G), which present the connections between all vertices in the network [4–6, 32]. Typically, rsfMRI matrices do not specify the direction of connectivity (*i.e.*, activity in area A drives activity in area B), therefore yielding undirected networks (Fig 1B and Fig 1F).

**Fig 1.**
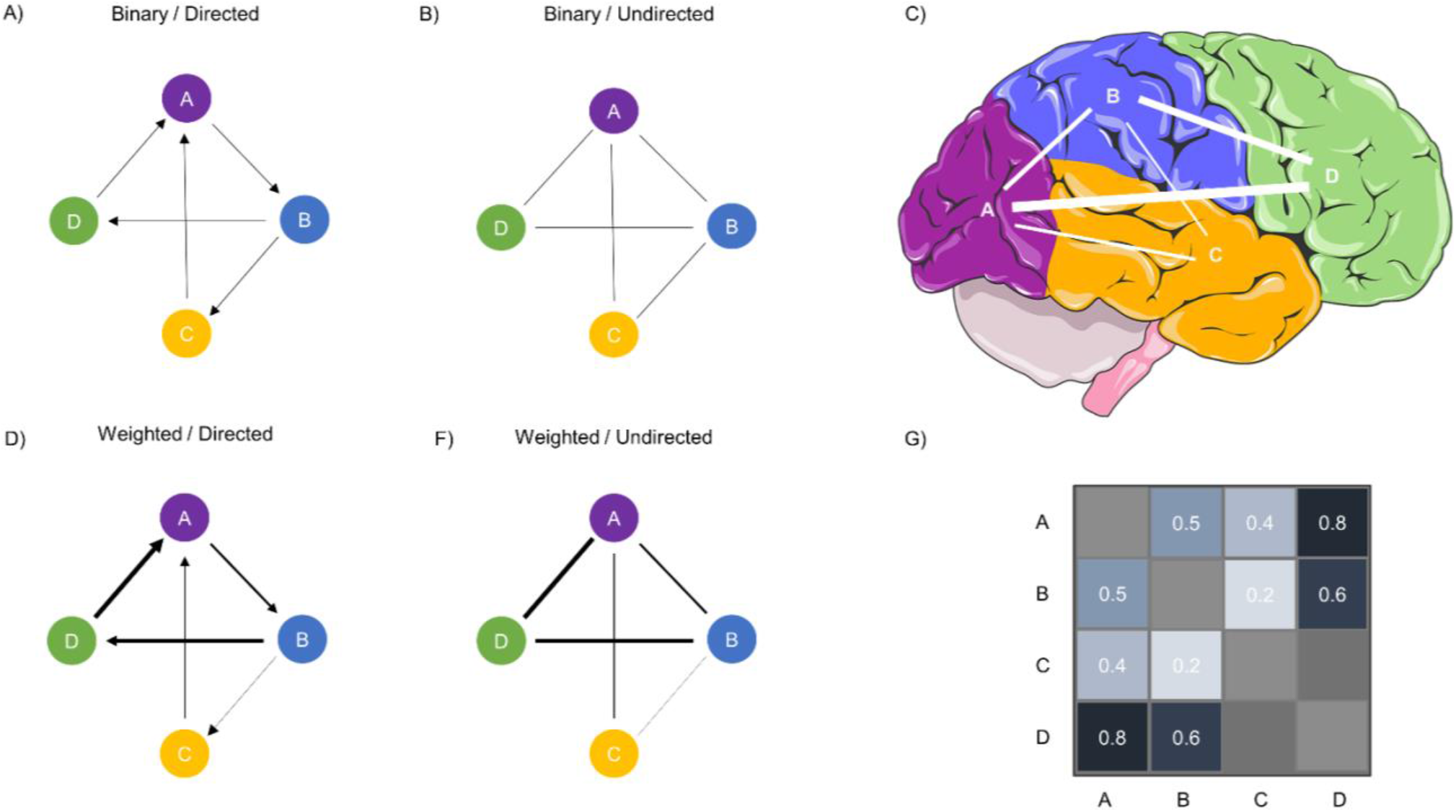
Types of networks. (A) A binary directed graph. (B) Binary, undirected graph. In binary graphs, the presence of a connection is signified by a 1 or 0 otherwise. (C) A representation of graph F as a network of brain areas. (D) A weighted, directed graph. (F) A weighted, undirected graph. In a weighted graph, the absolute strength of the connections is often represented by a number *w*, where 0 ≤ *w* ≤ 1. (G) A connectivity matrix of C and F. Source: Part of the image was obtained from smart.servier.com.

Before calculating any metrics on such matrices, several crucial factors must be considered when dealing with connectivity data [12, 33]. One critical decision is whether one wants to keep the information about edge weights. When the edges’ weights (*e.g.*, correlation values in rsfMRI connectivity) are maintained, the network will be weighted (Fig 1D and Fig 1F). Another approach is to use a threshold, *e.g.*, only keep and binarise the 20% strongest connections (Fig 1A and Fig 1B). There is currently no gold standard for the weighting issue in rsfMRI matrices [6, 33] and may also be dependent on the dataset or proposed analysis [34].

Another relevant discussion about rsfMRI matrices is the interpretation of negative weights or anticorrelations. The debate of what such negative correlations mean in neurophysiology is still going on [35]. Studies have suggested that they could be considered artefacts introduced by global signal regression or pre-processing methods, or simply by large phase differences in synchronised signals between brain areas [36, 37]. Nevertheless, a few authors have suggested that anticorrelations might carry biological meaning underlying long-range synchronisation and that in diseased states alterations in these negative correlations could indicate network reorganisation [35, 36]. Negative weights can be absolutised to keep the potential biological information they may carry. If one decides to discard them, it is crucial to keep in mind that some physiological information might be lost [6, 12, 35, 36].

In this tutorial, we will use an undirected, absolutised (positively) weighted matrix.

To follow the steps below, we assume that rsfMRI was already pre-processed and converted to a matrix according to some atlas. Steps and explanations on data pre-processing and atlas choices are beyond the scope of this paper, please see [38] for further information. Details on our Jupyter Notebook’s dataset pre-processing can be found in [39, 40].

#### Code example

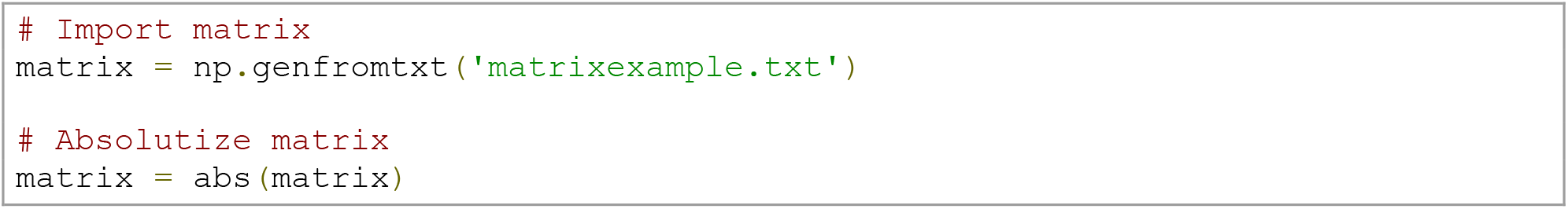

When working with fMRI brain network data, it is useful to generate some plots (*e.g.,* the heatmaps for matrix visualisation, and distribution plots of edge weights) to facilitate data exploration, comprehension and flag potential artefacts. In brain networks, we expect mostly weak edges and a smaller proportion of strong ones. When plotted as a probability density of log10, we expect the weight distribution to have a Gaussian-like form [6].

#### Code example

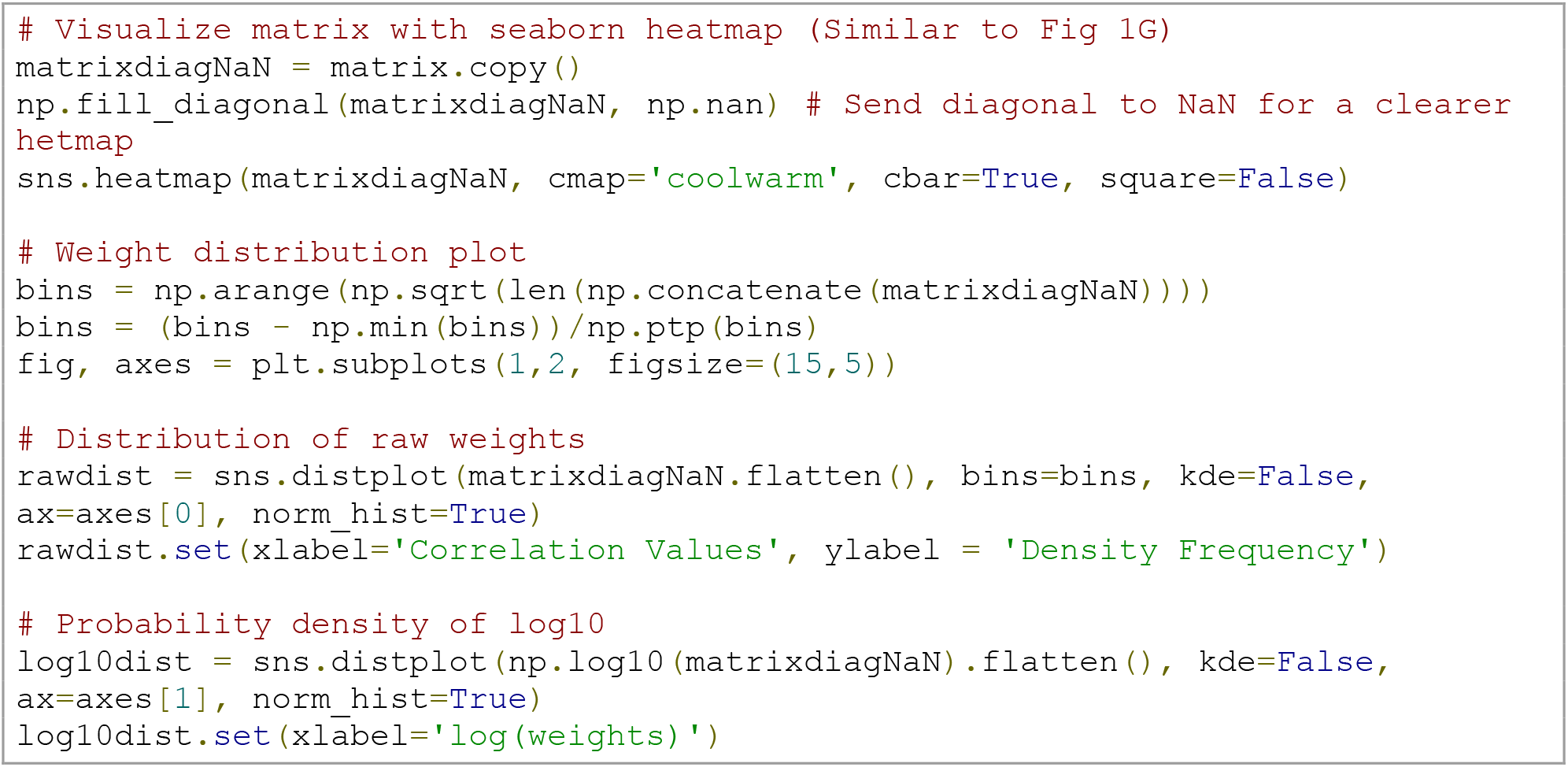

### Graph theory

Here, we will cover the most commonly used graph metrics in network neuroscience (see Fig 2), also in line with reference [6]. First, we need to create a graph object using the package *NetworkX* [41] and remove the self-loops (*i.e.,* the connectivity matrix’s diagonal).

**Fig 2.**
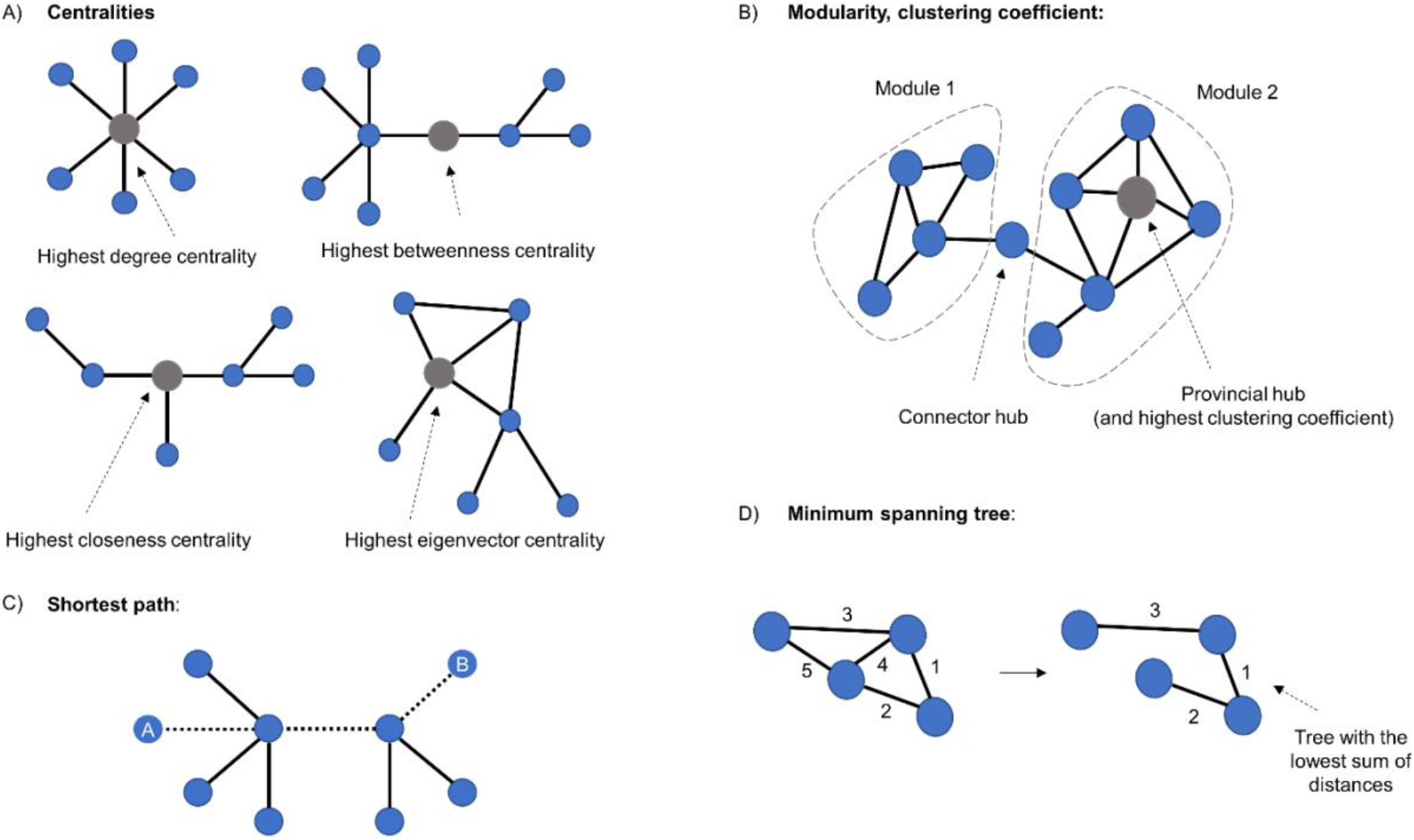
Graph theoretical metrics. (A) A representation of a graph indicating centralities. Highest degree centrality indicates the vertex with most connections. Highest betweenness centrality refers to the vertex with most short paths passing through it. Highest closeness centrality denotes the vertex that needs the least edges to reach all the other nodes. The highest eigenvector centrality is achieved by the vertex which is best connected to the rest of the network, considering the number of neighbours and how well connected they are. (B) Representation of modularity and clustering coefficient. The latter indicates the tendency for any two neighbours of a vertex to be directly connected to each other. (C) The shortest path between vertices A and B. (D) The minimum spanning tree is a subset of a graph’s edges, which does not contain cycles, and that has the lowest sum of distances.

#### Code example

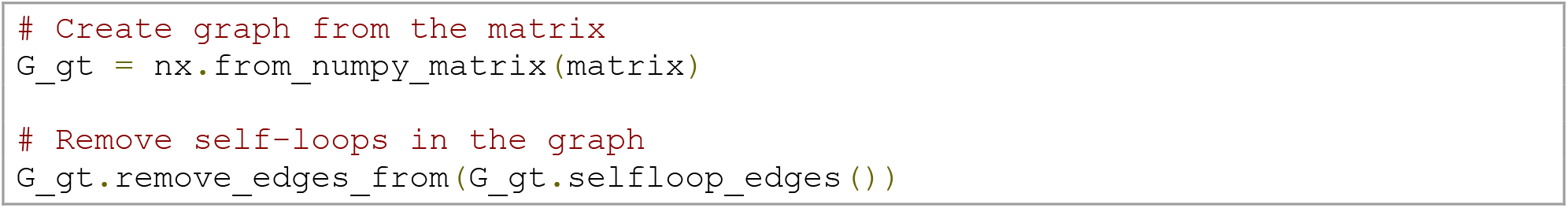

##### Degree

Vertex *degree* quantifies the total number of vertex connections in an undirected binary network [6]. In an undirected weighted network like our rsfMRI matrix, the vertex degree is analogous to the vertex *strength* (*i.e.,* the sum of all edges of a vertex) and equivalent to its degree centrality. This metric is one of the most fundamental metrics in network analysis and is a useful summary of how densely individual vertices are connected. It can be computed as the sum of edge weights of the neighbours of vertex *i*:

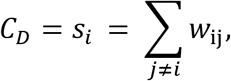

where *w*_*ij*_ is the weight of the edge linking vertices *i* and *j*.

#### Code example

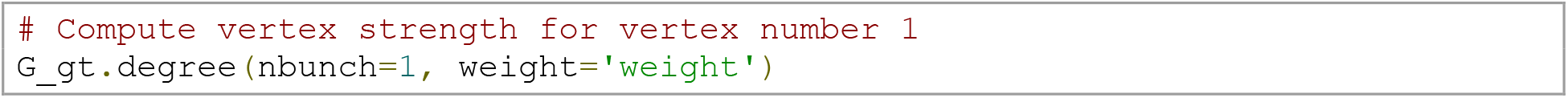

By removing the argument *weight* from the function, one can compute the degree of binarised networks where all edges are either 0 or 1 (useful if working with a sparse/not fully connected matrix). This change will give the vertex degree by calculating the number of edges adjacent to the vertex. One can also remove the specified vertex to estimate the degree/strength of all vertices. The *degree/strength distribution* allows us to scope the general network organisation in a single shot by displaying whether the network contains a few highly connected vertices, *i.e.,* hubs [12].

##### Path length

a. The *shortest path* is the path with least number of edges (or least total weight) between two vertices in a network. In a weighted graph, the shortest path is calculated by the minimum sum of the weights of edges between two vertices [6]. It is seen as a measure for understanding the efficiency of information diffusion in a network. Several algorithms can calculate path lengths, but Dijkstra’s algorithm [42] is one of the oldest and most well-known. An important detail is that this algorithm is only applicable to graphs with non-negative weights [42]. A pivotal point to keep in mind is that in the case of correlation matrices, such as rsfMRI data, the weights must be converted to ‘distance’ by computing the inverse of the original weight 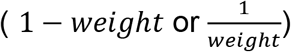; a higher correlation value represents a shorter distance [6]. This conversion is essential for all the following path-based metrics.
b. *Average path length (or characteristic path length)* is the average shortest path length for all possible pairs of vertices in a network. It is a global measure of information transport efficiency and integration in a network and is widely known due to the famous Watts-Strogatz model [14]. It can be computed as follows:

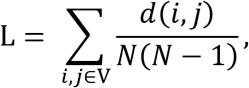

where *V* is a set of vertices, *d* (*i*, *j*) is the shortest path between vertices *i* and *j*, and *N* is the number of vertices in the network.

#### Code example

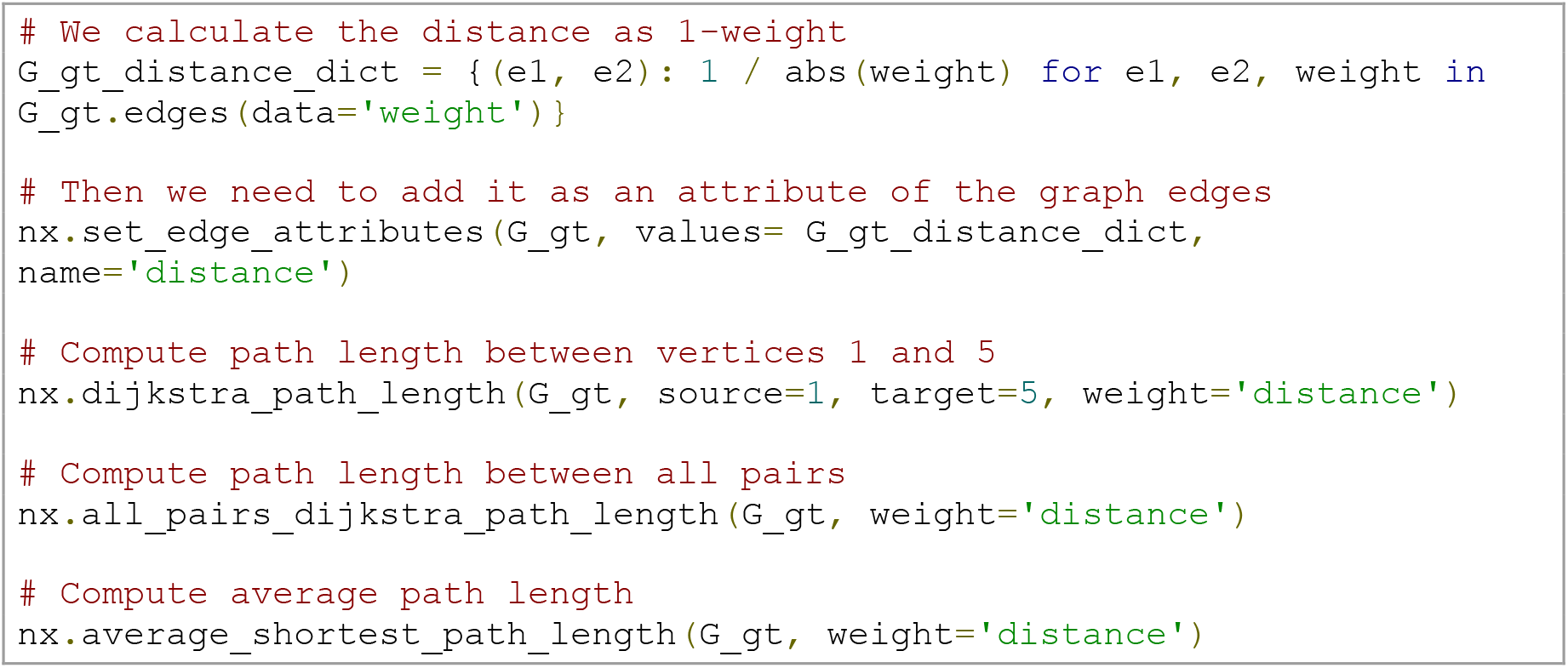

##### The clustering coefficient

The clustering coefficient assesses the tendency for any two neighbours of a vertex to be directly connected (or more strongly connected in the weighted case) to each other, and can also be termed cliquishness [12, 14]. This metric is also used for the computation of the small-worldness coefficient (ratio between the characteristic path length and the clustering coefficient relative to random networks) [14]. The formula can be defined as:

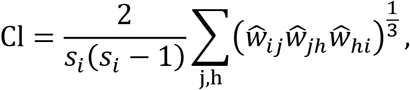

where *S*_*i*_ is the degree/strength of vertex *i*, and the edge weights’ are normalised by the maximum weight in the network, such that 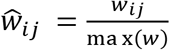.

#### Code example

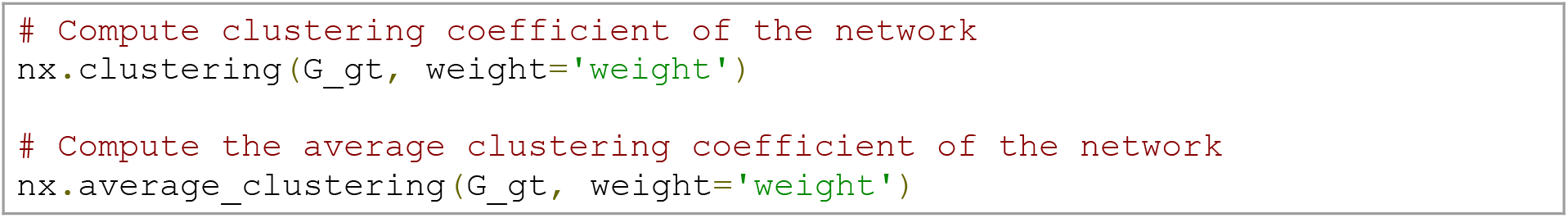

##### Centralities

a. *Eigenvector (degree-based) centrality* measures a vertex’s importance in a network while also considering how important its neighbours are [43]. Thus, it considers both the quantity and quality of a vertex’s connections. The eigenvector centrality can be computed as:

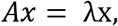

where A is the adjacency matrix, and x is an eigenvector of A with eigenvalue λ. We can also see the eigenvector centrality of a vertex *i* as the summed centrality of its neighbours:

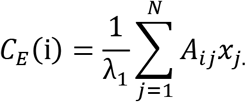 For weighted networks, certain conditions apply. According to the Perron-Frobenius theorem, the adjacency matrix’s largest eigenvalue must be unique and positive, which is guaranteed only for matrices with positive values [6, 44].
b. *Closeness (shortest path-based) centrality* is a measure of how closely or’ directly’ connected a vertex is to the rest of the network. If the vertex is the closest to every other element in the network, it has the potential to spread information fast and efficiently [6]. Formally, the closeness centrality of a vertex *i* is the inverse of its average shortest path length (*N.B.* weights need to be converted to distances) to all N-1 other vertices:

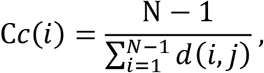

where d(*i*, *j*) is the shortest-path distance between *i* and *j*, and *N* is the number of vertices in the graph. In weighted networks, closeness centrality can be estimated by considering the summed weight of the shortest paths according to Dijkstra’s algorithm [42].
c. *Betweenness (shortest path-based) centrality* is the proportion of all vertex-pairs shortest paths in a network that pass through a particular vertex [44, 45]. It is used to understand the importance of vertices in the overall flow of information in a network. To compute the betweenness centrality of a vertex *i*, one has to calculate the proportion of shortest paths between two vertices, *e.g.*, *i*, *j*, that pass through vertex *h*:

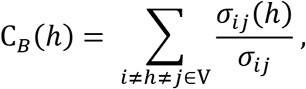

where *V* is a set of vertices, *σ*_*ij*_ is the total number of shortest paths between *i* and *j*, and *σ*_*ij*_(*h*) is the number of those paths that pass through *h*. For weighted graphs, edges must be greater than zero, and the metric considers the sum of the weights [6]. Again, it is necessary to use the distance when using this shortest path-based metric. This formula can also be normalised by putting 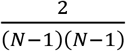 in front of the sum (N being the number of vertices).

#### Code example

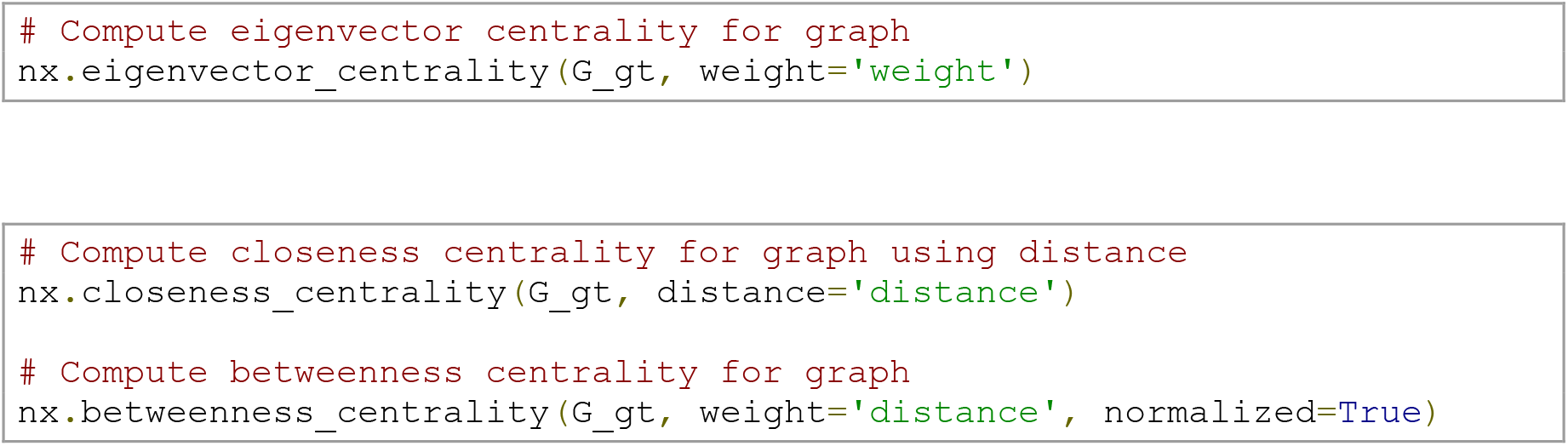

##### The minimum spanning tree

The minimum spanning tree is the backbone of a network, *i.e.*, the minimum set of edges necessary to ensure that paths exist between all vertices without forming cycles [46, 47]. A few main algorithms are used to build the spanning tree, with Kruskal’s algorithm being implemented in *NetworkX* [48]. Briefly, this algorithm ranks the distances between vertices, adds the ones with the smallest distance first, and adding edge by edge it checks if cycles are formed or not. The algorithm will not add an edge that results in the formation of a cycle.

#### Code example

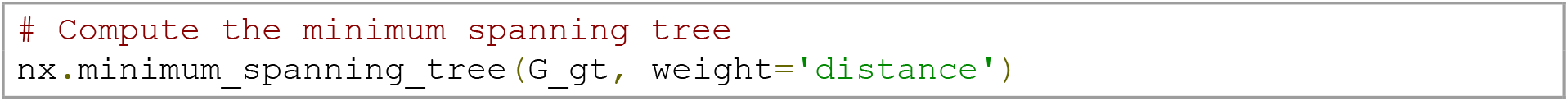

##### Modularity

Modularity states how divisible a network is into different modules (or communities) and the identification of the modules is performed by the community detection algorithm [6, 8, 49]. Here, we will use the Louvain algorithm [50] as recommended by [6]. It works in a two-step iterative manner, first looking for communities by optimising modularity locally and then concatenating vertices that belong to the same module [50].

#### Code example

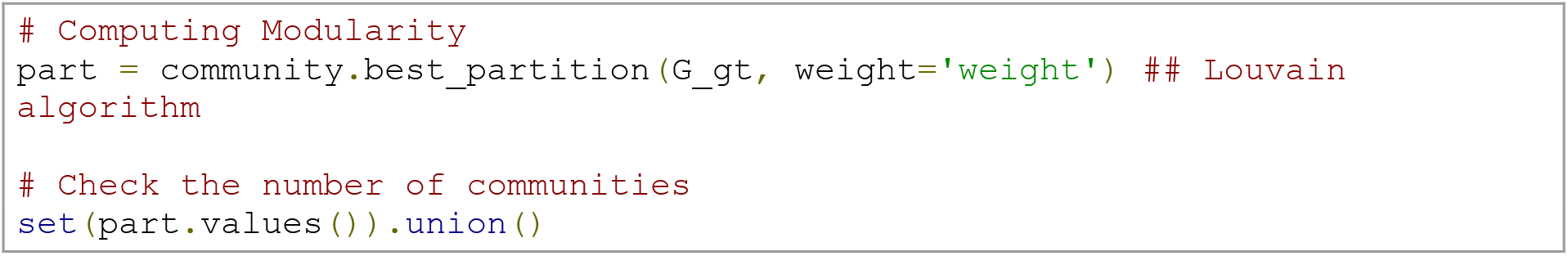

### Topological data analysis

In this section, we will use TDA on our rsfMRI adjacency matrices. TDA can identify different characteristics of a network by addressing the high-order structure of a network beyond pair-wise connections as used in graph theory [30, 51, 52]. TDA generally uses topology and geometry methods to study the shape of the data [53]. A core success of TDA is the ability to provide robust results when compared with alternative methods, even if the data are noisy [18, 23]. One of the benefits of using TDA in network neuroscience is the possibility of finding global properties of a network that are preserved regardless of the way we represent the network [25], as we will illustrate below. Those properties are the so-called topological invariants.

We will cover some fundamental TDA concepts: filtration, simplicial complexes, Euler characteristic, phase-transitions, Betti numbers, curvature, and persistent homology. A summary can be found in Fig 3.

**Fig 3.**
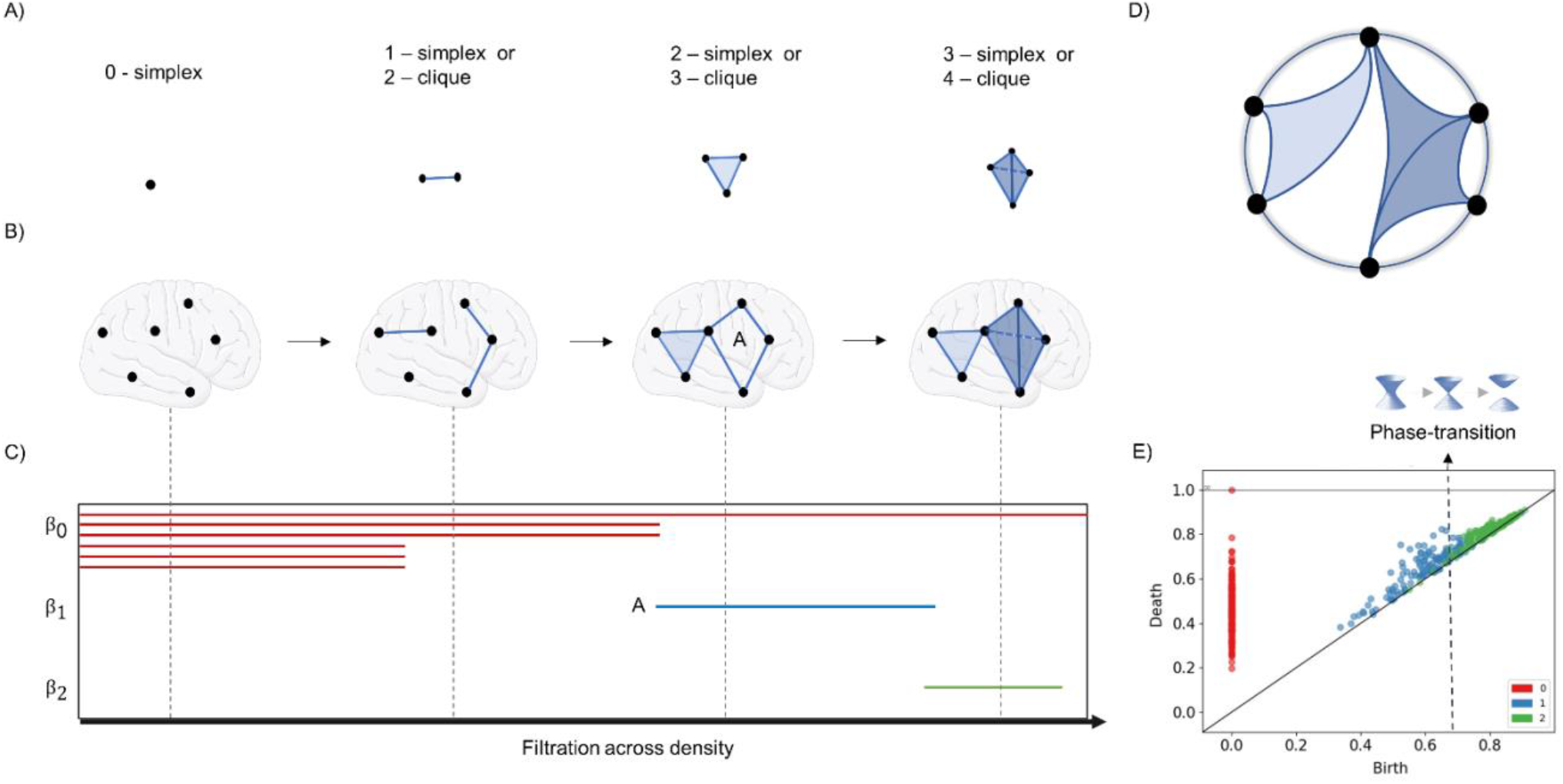
Topological data analysis. (A) Illustration of simplexes. (B) Representation of simplexes/cliques of different order being formed in the brain across the filtration process. (C) Barcode respective to panel B, representing the filtration across distances (*i.e.*, the inverse of weights in a correlation matrix). Line A represents cycle A in B. *β*_0_, *β*_1_, and *β*_3_ indicate the homology groups. (*β*_0_ =connected components, *β*_1_ = one-dimensional holes, *β*_2_ = 2-dimensional holes). (D) Circular projection of how the brain would be connected. (E) Persistence diagram (or Birth/Death plot) obtained from real rsfMRI brain data. In this plot, it is also possible to identify a phase transition between *β*_1_and *β*_2_.

#### The basis: the adjacency matrix and filtration

As indicated in the earlier section on graph theory, there is no consensus on the necessity or level of thresholding performed on rsfMRI-based adjacency matrices. However, TDA overcomes this problem by investigating connectivity over all possible thresholds in a network. This process of investigating network properties looking for all possible thresholds instead of choosing a fixed one is called *filtration* (Fig 3B and S1 Video). It consists of changing the threshold, *e.g.,* the density *d* of the network, from 0 ≤ *d* ≤ 1. This yields a nested sequence of networks, in which increasing *d* leads to a more densely connected network. Note, however, that the notion of filtration is not unique to TDA and has also been applied in graph-theoretical work [26, 54].

### Code example

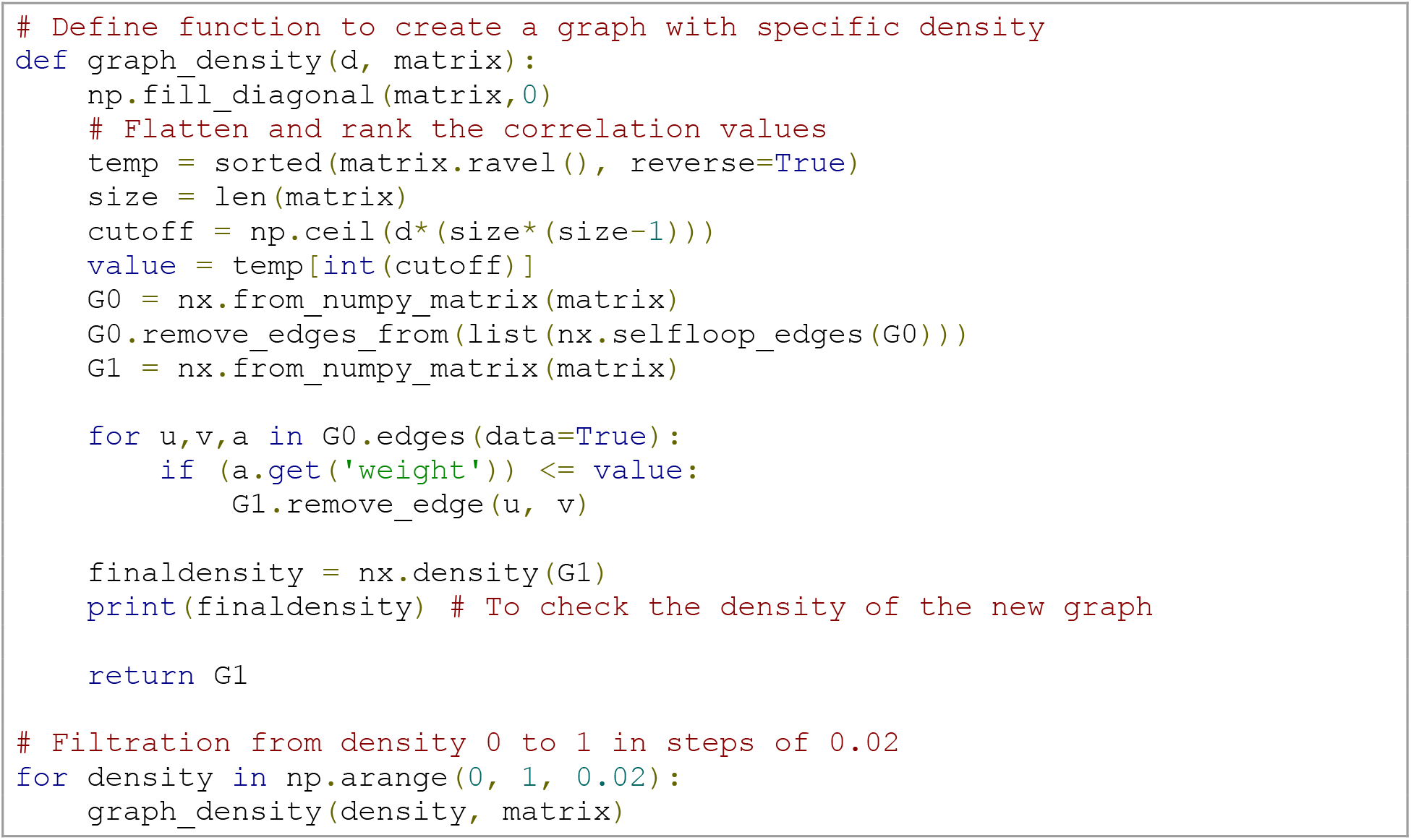

#### Simplicial complexes

In TDA, we consider that the network as a multidimensional structure called the simplicial complex. Such a network is not only made up of the set of vertices (0-simplex) and edges (1-simplex) but also of triangles (2-simplex), tetrahedrons (3-simplex), and higher *k*-dimensional structures (Fig 3A). In short, a *k*-simplex is an object in *k-*dimensions and, in our work, is formed by a subset of *k+1* vertices of the network (Fig 4).

**Fig 4.**
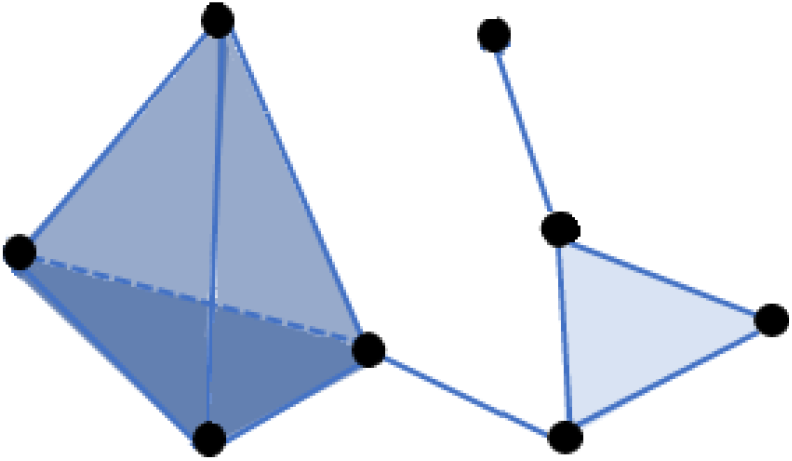
Simplicial complex. An example of a simplicial complex composed of eight vertices (0-simplexes), 11 edges (1-simplexes), five triangles (2-simplexes), one tetrahedron (3-simplexes).

We can encode a network into a simplicial complex in several ways [55–57]. However, here, we will focus on building a simplicial complex only from the brain network’s cliques, *i.e.*, we will create the so-called clique complex of a brain network. In a network, a k-clique is a subset of the network with *k* all-to-all connected nodes. 0-clique corresponds to the empty set, 1-cliques correspond to nodes, 2-cliques to links, 3-cliques to triangles, and so on. In the clique complex, each *k+1* clique is associated with a *k*-simplex. This choice for creating simplexes from cliques has the advantage that we can still use pair-wise signal processing to create a simplicial complex from brain networks, such as in [31]. It is essential to mention that other strategies to build simplicial complexes beyond pair-wise signal processing are still under development, such as applications using multivariate information theory together with tools from algebraic topology [58–63].

In our Jupyter Notebook, we provide the code to visualise the clique complex developed in [64]. To create the 3-D plots, we used mesh algorithms available in *Plotly* [65], together with a mesh surface of the entire brain available in [66, 67]. In Fig 5, we display an example of 3-D visualisation of 3-cliques in the HCP data. When we increase the filtration density *d*, we obtain more connections, and more 3-cliques arise. In Fig 5, only 3-cliques are shown; however, the same can be done for higher-dimensional cliques like a tetrahedron, *et cetera*. In the S1 Video, we show a filtration in a functional brain network up to 4-cliques. In the Jupyter Notebook, we are also able to visualise the clique complex at arbitrary sizes, up to computational limits. The computation is not shown here as code blocks due to its size and complexity (see the Jupyter Notebook).

**Fig 5.**
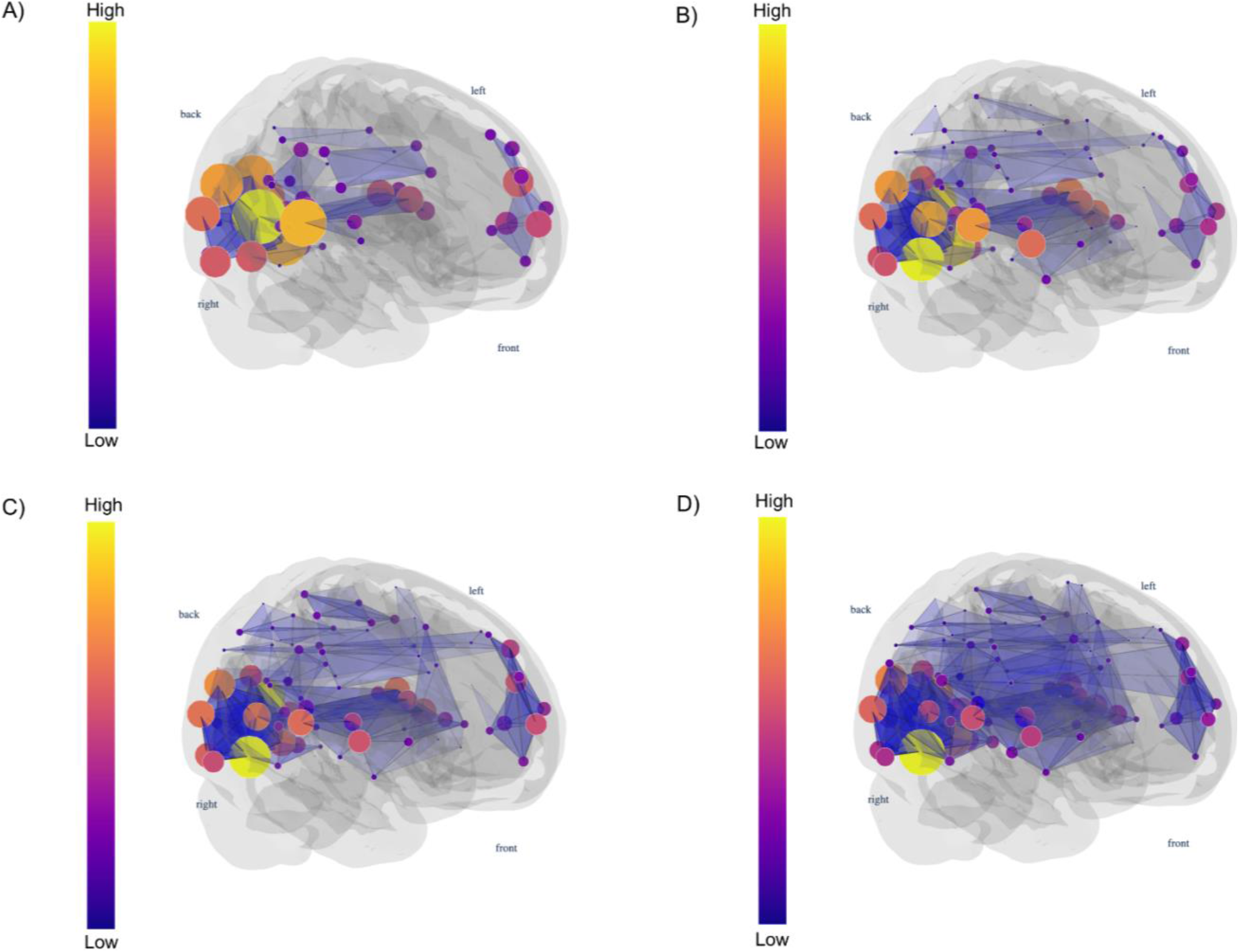
Simplex 3-D visualisation. Here we visualise the rising number of 3-cliques (triangles) in a functional brain network as we increase the edge density *d* (0.01, 0.015, 0.02, and 0.025, from A to D). For higher densities, we have a more significant number of 3-cliques compared to smaller densities. The vertex colour indicates the clique participation rank.

#### The Euler characteristic

The Euler characteristic is one example of topological invariants: the network properties that do not depend on a specific graph representation. We first introduce the Euler characteristic for polyhedra, as illustrated in Fig. 6. Later, we translate this concept to brain networks. In 3-D convex polyhedra (for example a cube, a tetrahedron, *et cetera,* see Fig. 6), the Euler characteristic is defined as the numbers of vertices minus edges plus faces of the considered polyhedra. For convex polyhedra without cavities (holes in its shape), which are isomorphous to the sphere, the Euler characteristic is always two, as you can see in Fig 6. If we take the cube and make a cavity, the Euler drops to zero as it is in the torus. If we make two cavities in a polyhedral (as in the bitorus), the Euler drops to minus two (Fig 7). We can understand that the Euler characteristic tells us something about a polyhedron’s topology and its analogous surface. In other words, if we have a surface and we make a discrete representation of it (*e.g.*, a surface triangulation), its Euler characteristic will always be the same, regardless of the way we do it.

**Fig 6.**
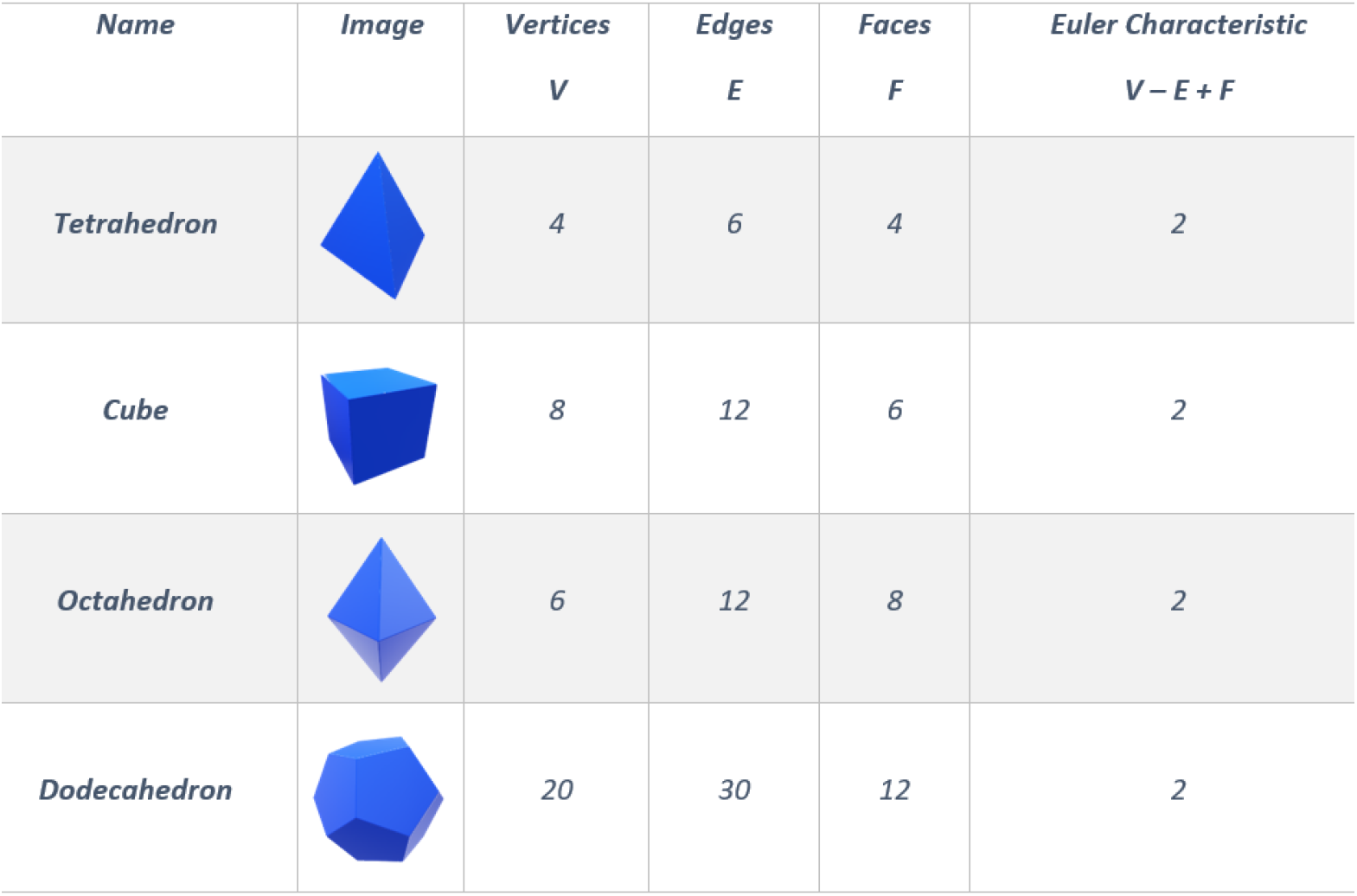
Euler characteristic in convex polyhedra. Note that there are no cavities in their shapes for convex polyhedra, and the Euler characteristic is always equal to two.

**Fig 7.**
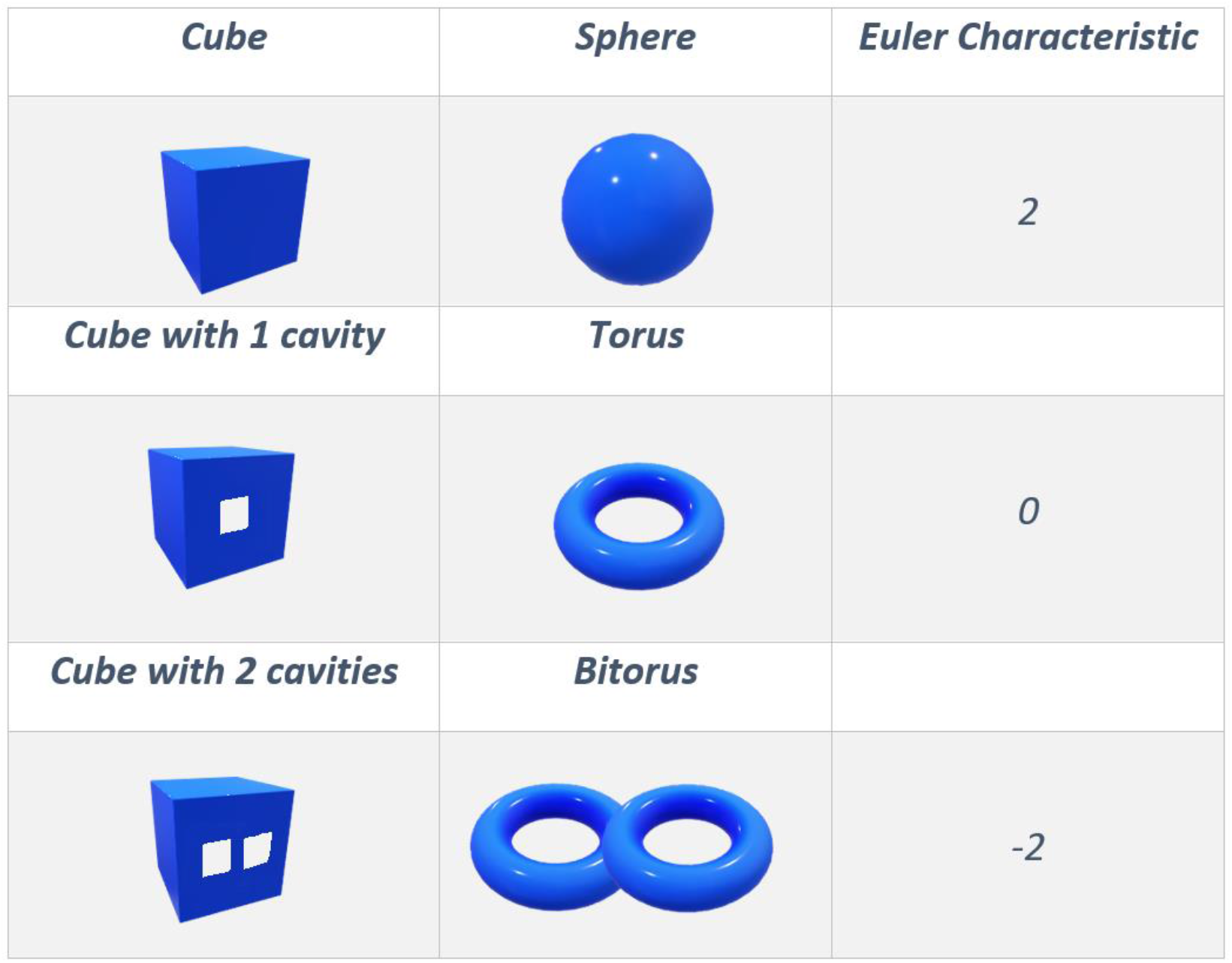
The Euler characteristic in polyhedra with cavities. The Euler characteristic of a cube with a cavity is equal to zero, just as the torus. This value drops to minus two if we have two cavities in the cube, just like a bitorus.

We can now generalise the definition of Euler characteristic to simplicial complex in any dimension. Thus, the high dimensional version of the Euler characteristic is expressed by the alternate sum of the numbers *Cl*_*k*_ (*d*) of the *k*-cliques (which are (*k-1)*-simplexes) present in the network’s simplicial complex for a given value of the density threshold *d*.

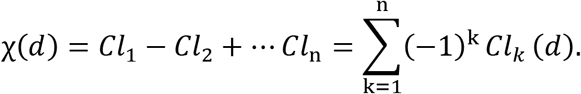

### Code example

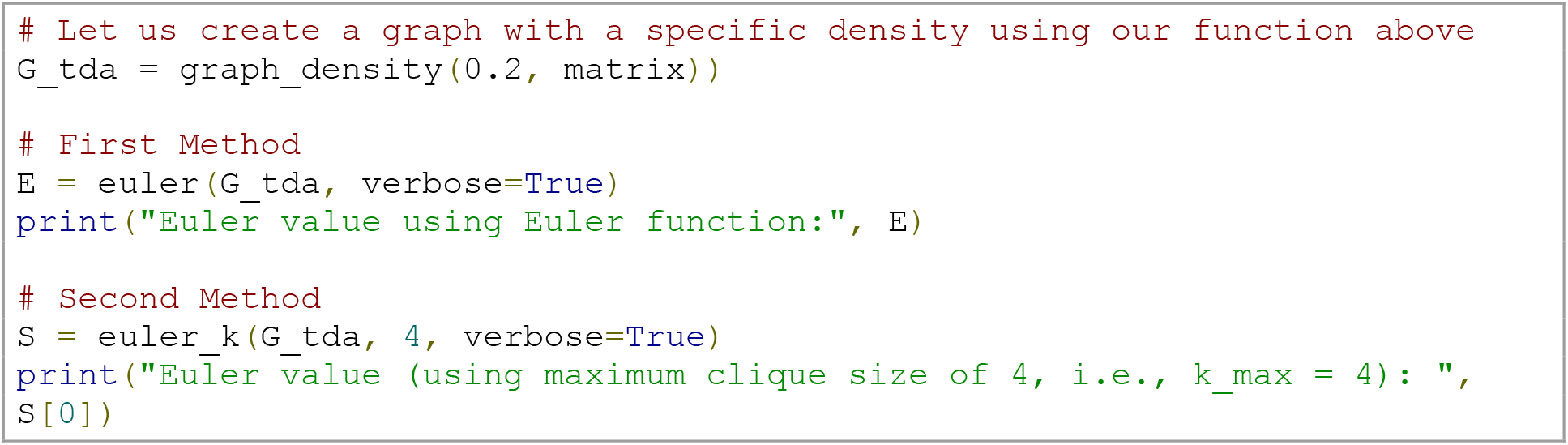

Note that the clique algorithm, which is the primary function used in our code (*euler* - S1 Code Block), is an NP-complete problem, which is computationally expensive for large and/or dense networks, regardless of the way you implement it [68]. An alternative is to fix an upper bound for the cliques’ size [68, 69]. Therefore, the second function (*euler_k* - S1 Code Block) allows the user to constrain the maximum size of the cliques we are looking for. This means that we are fixing the dimension *k* of our simplicial complex and ignoring simplexes of dimension greater than *k*.

#### Topological phase transitions

Phase transitions can provide insight into the proprieties of a ‘material’. For example, water is known for becoming steam at 100°C. Similarly, by using TDA when comparing a patient and healthy population, one could identify these populations’ properties by studying the topological phase transition profile for each group. This strategy has already been applied for investigating group differences between controls and glioma brain networks [64], and typically developing children and children with attention-deficit/hyperactivity disorder [26]. In other fields, topological phase transitions were also investigated in the *S. cerevisiae* and *C. elegans protein interaction networks,* reionization processes, and evolving coauthorship networks [70–72].

To investigate topological phase transitions in brain networks, we first need to visualise the *Euler entropy* (Fig 3 in [73]):

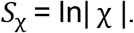

When χ = 0 for a given value of the density of the network, the Euler entropy is singular, *S*_χ_ → ∞. Under specific hypotheses, a topological phase transition in a complex network occurs when the Euler characteristic is null [64]. This statement finds support in the behaviour of *S*_χ_ at the thermodynamic phase transitions across various physical systems [73]. In network theory, the Giant component transition is associated with network changes, from smaller connected clusters to the emergence of Giant ones [74]. Theoretically, topological phase transitions are related to the extension of the Giant component transition for simplicial complexes [75]. Based on numerical simulations, it was also conjectured that the longest cycle is born in the phase transition vicinity [76, 77]. Phase-transitions can also be visualised in Birth/Death plots (Fig 3E) which will be discussed later in the Persistent Homology section.

#### Betti numbers

Another set of topological invariants are the Betti numbers (*β*). Given that a simplicial complex is a high-dimensional structure, *β*_*k*_ counts the number of *k*-dimensional holes in the simplicial complex. These are topological invariants that correspond, for each *k* ≥0, to the number of linearly independent *k*-dimensional holes in the simplicial complex [16].

In Fig 8, we show the representation of the *k*-dimensional holes. We give one example for each dimension. In a simplicial complex, there can be many of these *k*-holes and counting them provide the Betti number *β*, *e.g.,* if *β*_2_ is equal to five, there are 5 two dimensional holes.

**Fig 8.**
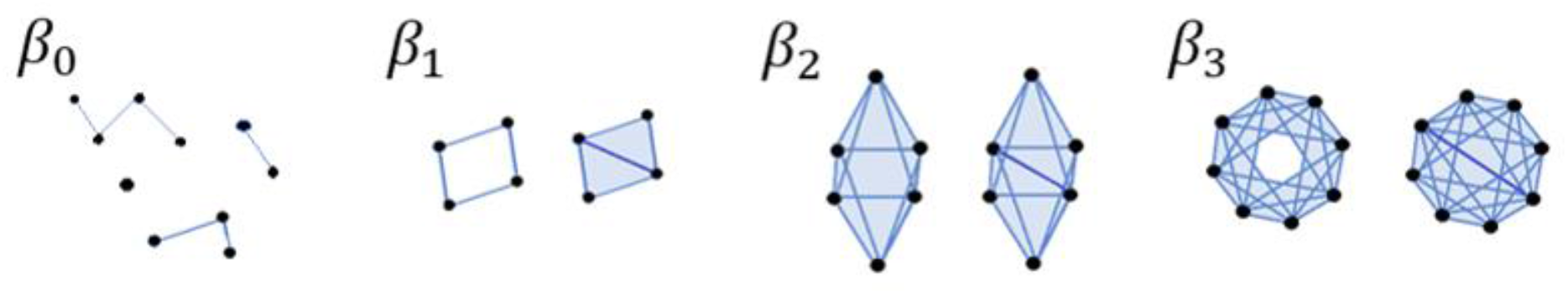
Betti numbers and examples of each *k*-dimensional hole. *β*_0_ is the number of connected components or zero-dimensional holes. *β*_1_ is the number of one-dimensional holes (loops). *β*_2_ is the number of two-dimensional holes (voids). *β*_3_ is the number of 3-D holes. For *β*_1_, *β*_2_, *β*_3_ only the left figure of each pair represents the k-dimensional hole. In the right figure, a connection is added, and so the k-hole is lost: the right figure of each pair no longer represent a *β* hole. For *β*_0_ the number of connected components is the number of separate clusters we have in the figure; therefore, we should consider the figure as a whole (in the case represented here we have four connected components).

### Code example

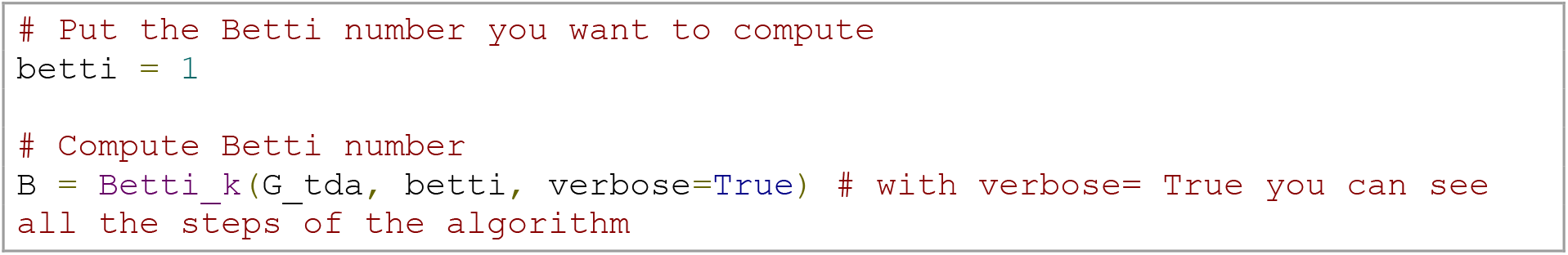

Notice that for higher k or dense simplicial complexes, the calculation of the *β* becomes computationally expensive.

There are ways to estimate the *β* of a simplicial complex without calculating it directly. It is known that the *β* relate to Euler characteristics and phase transitions. The Euler characteristics of a simplicial complex can also be computed using the *β* via the following formula [56]:

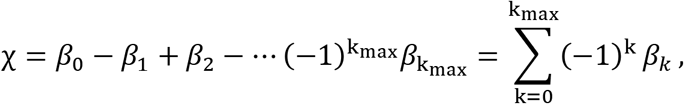

where *k*_max is the maximum dimension that we are computing the cycles.

Furthermore, topological phase transitions are also defined in terms of the *β* of a simplicial complex [78]. We know that β_0_ counts the number of the connected components of a simplicial complex. If we compute the Betti curves as a function of probability in stochastic models, each Betti curve passes through two distinct phases, in a narrow interval: one when it first emerges, and the other when it vanishes [75]. That means that, under similar assumptions as in theoretical models, if the *β* distribution is unimodal, increasing the density of edges of a brain network will lead to the appearance of *β* of a higher order. In contrast, smaller Betti numbers will disappear at the vicinity of a topological phase transition.

In Figure 9, we illustrate this property on simplicial complexes obtained from random networks. As the probability increases and so the density of the network is higher, we find a sequence of dominant *β*_*k*_ starting from *k*=0, that change (*i.e.*, *k* is incremented by one unity) every time a topological phase transition occurs. While the singularities of the Euler entropy *S*_χ_ determine the transitions’ location, the crossover of the *β* characterises which kind of multidimensional hole prevails in each topological phase of the filtration.

**Fig 9.**
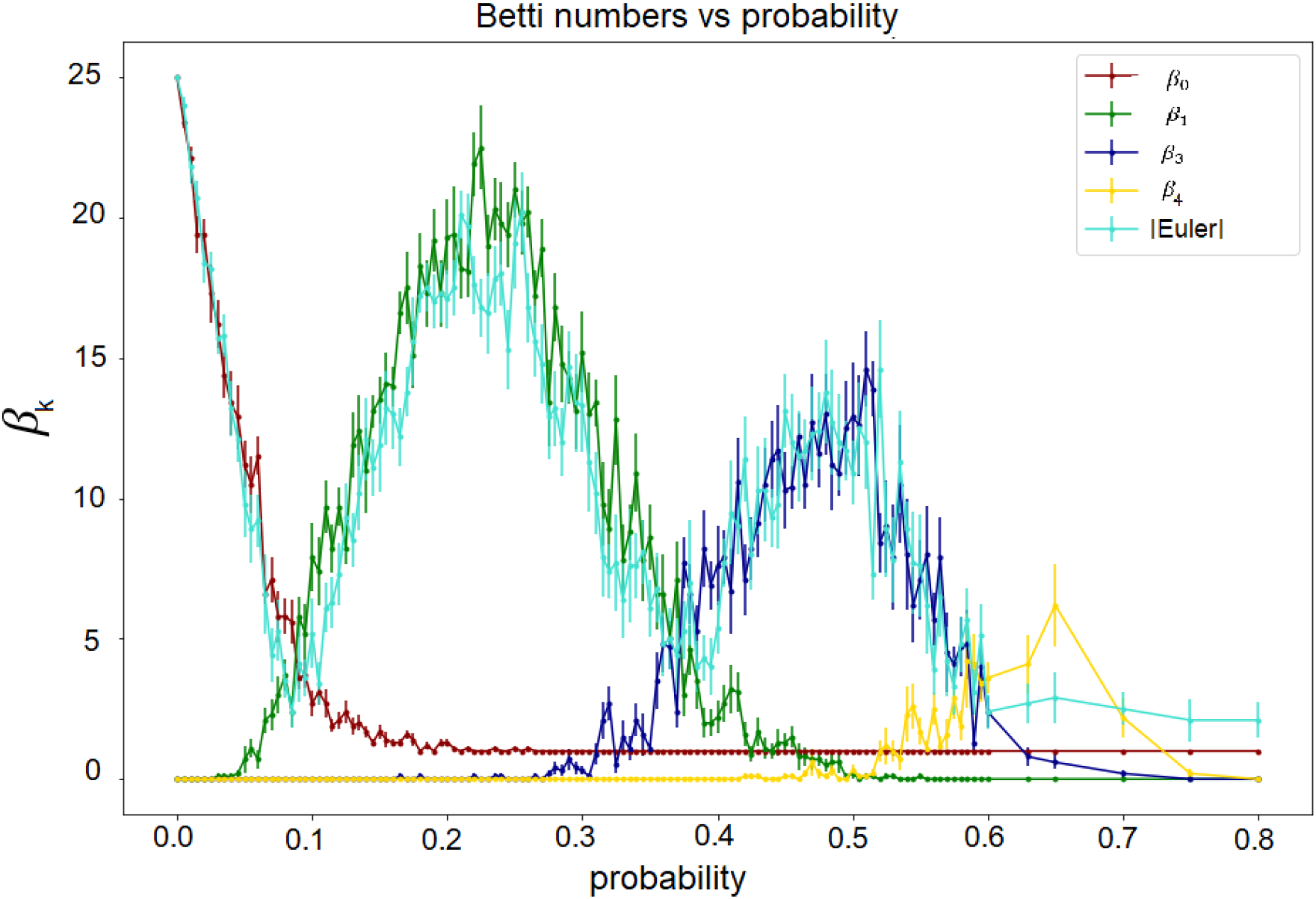
Betti number and Euler characteristic approximation. We create a random network for each probability of connection between vertices, and we compute the *β* and the absolute value of the Euler characteristic. We repeated the experiment 10 times and calculated the mean curves with errors. In this plot, we only show the mean (with errors) of the ten experiments for the Euler and Betti curves. We notice that the absolute value of the Euler characteristic is a good approximation of *β*.

#### Curvature

Curvature is a TDA metric that can link the global network properties described above to local features [64, 79, 80]. When working with brain network data, this will allow us not only to compute topological invariants for the whole-brain set of vertices but also to understand the contribution of specific individual nodal, or subnetwork, geometric proprieties to global properties of the brain network.

Several approaches to defining a curvature for networks are available [19], including some already used in neuroscientific investigations. We will illustrate the curvature approach linked to topological phase transitions, previously introduced for complex systems in [80–82].

To compute the curvature (S3 Code Block), filtration is used to calculate the clique participation rank (*i.e.*, the number of *k*-cliques in which a vertex *i* participates for density *d*) [3], which we denote here by *Cl*_*ik*_ (*d*). The curvature of the vertex based on the participation rank is then defined as:

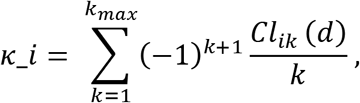

where *Cl*_*ik*_ = 1 since each vertex *i* participates in a single 1-clique (the vertex itself), and *k*_*max*_ the maximum number of vertices that are all-to-all connected in the network.

To link this nodal curvature to the network’s global properties, we use the Gauss-Bonnet theorem for networks, through which one can connect a local curvature of a network and its Euler characteristic. Conversely, by summing up all the curvatures of the network across different thresholds, one can reach the alternate sum of the numbers *Cl*_*k*_ of *k*-cliques (a subgraph with *k* all-to-all connected vertices) present in the simplicial complex of the network for a given density threshold d ∈ [0, 1], according to the following equation:

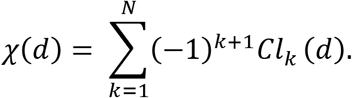

By doing so, we also write the Euler characteristics as a sum of the curvature of all network vertices, *i.e.*,

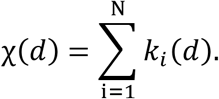

We illustrate the curvature distribution for a functional brain network for densities before and after the transition in Fig 10.

**Fig 10.**
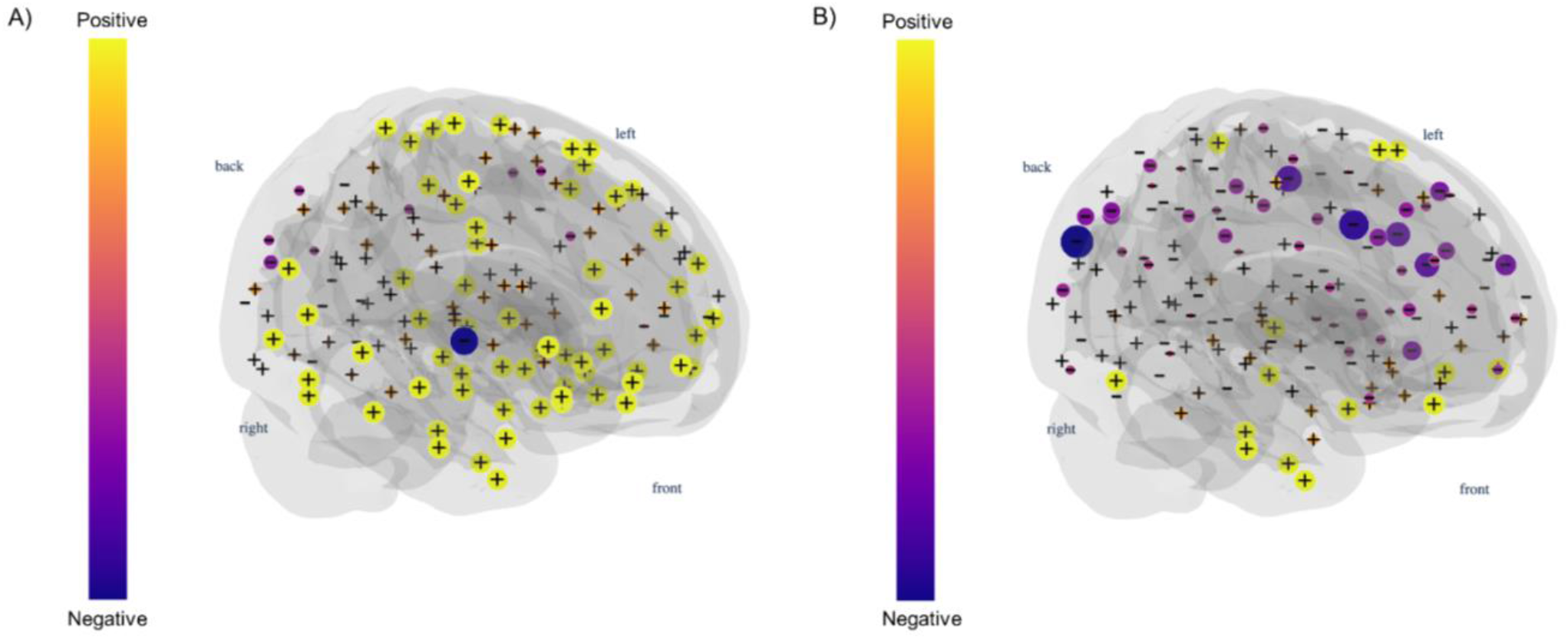
Curvature 3-D plot. Distribution of curvatures in a functional brain network for densities 0.01 (A) and 0.03 (B) after the first topological phase transition. The sum of curvature over all vertices is equal to the Euler characteristic.

#### Persistent homology

Homology is a topology branch that investigates objects’ shape by studying their holes (or cycles). Persistent homology tracks the emergence of cycles across the evolving simplicial complexes during filtration, allowing us to recognise whether there were homology classes that “persisted” for many filtrations (time here meaning the threshold gap between the birth and death of a cycle) [24, 28]. Importantly, to compute persistent homology, we need to work with a distance matrix, which is the first step in the code below. We can then calculate the simplicial complex’s persistence and plot it as a barcode or as a persistence diagram (Fig 3C and E). Here we used the *Gudhi* package for the implementation of those steps [83]. The topological phase transitions in complex networks [64, 70], which can also be identified between the changes in the dimensionality of the birth/death graphs mentioned above (Fig 3E).

### Code example

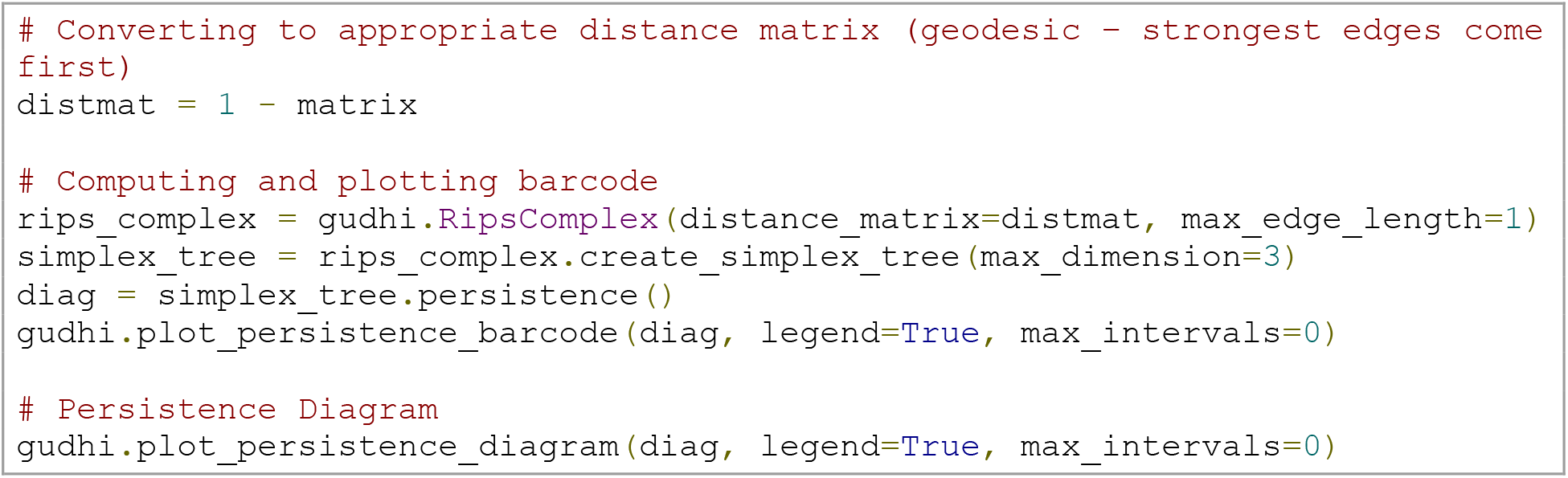

## Discussion

This tutorial has explained some of the main metrics related to two network neuroscience branches - graph theory and TDA-, providing short theoretical backgrounds and code examples accompanied by a publicly available Jupyter Notebook. We innovate by combining hands-on explanations with ready-to-use codes of these subfields and visualisations of simplicial complexes in the brain, thereby hopefully lowering the high threshold necessary for neuroscientists to get acquainted with these new analysis methods, particularly for rsfMRI data. Here, we also innovate by providing realistic visualization of higher-order simplices in brain networks.

Notably, there are limitations and other relevant points that should be kept in mind when working with these metrics. Firstly, it is a common practice in network neuroscience to use null models for comparison with real data. The idea is to show that the results are different from what one would obtain by chance (or randomly). The generation and comparison with null models must be performed differently for graph theory and TDA, and it is crucial to define what propriety should be kept constant (*e.g.*, the density of the network, or degree distribution). Nevertheless, the computation and discussion of null models are beyond this tutorial’s scope and would be an article in itself. A more in-depth discussion of null models in graph theory can be found in [6]. Please see section 4 of [42], and [72] for null models in simplicial complexes (and see Table 1 for further resources).

Moreover, it is crucial to appreciate limitations in interpretation when using these metrics in connectivity-based data. Considering that rsfMRI data is often calculated as a temporal correlation between time series using Pearson’s correlation coefficient, a bias on the number of triangles can emerge. For example, suppose areas A and B, and areas C and B are communicating and thus correlated. In that case, a correlation will be present between A and C, even if there would be no actual communication between these vertices [84]. This can affect graph-theoretical metrics such as the clustering coefficient, with networks based on this statistical method being automatically more clustered than random models, and TDA metrics, where the impact depends on how high-order interactions are defined. The proper way to determine and infer high-order interactions in the brain is an ongoing challenge in network neuroscience. Here we simplified our approach using the cliques of a network to define our simplicial complex. For those interested in a more in-depth discussion on the topic, we recommend section 1 and 3 of chapters 7 and 10, respectively, in [6].

The use of weighted matrices can also come with caveats. As mentioned above, various metrics use the sum of weights to compute final nodal values. From that, multiple edges with low weights might have a final sum equal to few edges with higher weights. How to deal with this limitation and distinguish between these cases is still under discussion. A possible solution was proposed by [85], in which the addition of a tunable parameter in the computation of centralities can allow the researcher to include the number of edges in the total sum, not only the sum of the weights.

Concerning TDA, it is essential to think about limitations in its use due to computational power. The computation of cliques falls in the clique-problem, an NP (nonpolynomial time) problem, thus listing cliques may require exponential time as the size of the cliques or networks grows [69]. For example, if the matrix to be analysed has 60 vertices with a maximum clique size of 23, this will correspond to 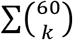 for *k* ∈ {0,…, 23} cliques, resulting in an enormous taimeoutontcoofmpute all cliques. What we can do for practical applications is to limit the clique size that can be reached by the algorithm, which determines the dimension of the simplicial complex in which the brain network is represented. This arbitrary constraint implies a theoretical simplification, limiting the space or the dimensionality in which we would analyse brain data. Another issue is that, to finish TDA computations in a realistic timeframe, the researcher might need to establish a maximal threshold/density for convergence even after reducing the maximal clique size. Even though TDA approaches lead to substantial improvements in network science; apart from applications using the Mapper algorithm [86], the limitations mentioned above contribute to losing information on the data’s shape [87].

Furthermore, given the early stage of TDA approaches in clinical network neuroscience, it is relevant to recognise that the neurobiological meaning of the metrics mentioned here is still not well defined. Further studies contrasting different neuroscientific techniques with TDA must be done to explain in neurobiological level what a topological metrics represent and how they correlate with brain functioning. However, it is already possible to use these metrics to differentiate groups [26, 64], and plausible to assume that the interpretation of some classical metrics could be extrapolated to higher orders interactions. For example, the concept of the centralities using pair-wise interactions is used to understand node importance and hubs, the same, in theory, could be applied to the relationships between 3 or more vertices by extending the definition of centrality from networks to simplicial complexes, as done in [88, 89].

Lastly, we would like to briefly mention more general problems in network neuroscience and brain imaging. Before embarking on the application of graph theoretical or topological data analysis, one should be aware of frequent arbitrary decisions such as defining thresholds, using binary or weighted matrices, and controlling for density. Besides, one should think about the differences that arise from using particular atlases and parcellations and their influence on the findings [6, 8, 26, 90–93]. All these factors can impact how credible and reproducible the field of network neuroscience will be, inevitably influencing how appealing the metrics’ use might be to clinical practice [91].

## Conclusion

Network neuroscience is pivotal in the understanding of brain organisation and function. Graph theory has been the most utilised framework so far, but as the field of network neuroscience expands, newer methods such as TDA are starting to take part in the investigation. To further improve the field, especially in clinical network neuroscience, it is imperative to make the computation of the developed metrics accessible, easy to comprehend, visualize, and efficient. Moreover, researchers must be aware of the crucial decisions that one must make when executing data analysis and how these can affect studies’ results and reproducibility. We hope to have facilitated the comprehension of some aspects of network and topological neuroscience, the computation and visualization of some of its metrics. As a final reminder, we would again suggest that the reader explores our table of resources, and the Jupyter Notebook developed by our team.

## Supporting information

Supplemental Code blocks 1-3

Sup. video: Filtration in a brain network

## Supporting information

**S1 Video. 3-D filtration of networks with 2,3,4 and 5-cliques.**

**S1 Code block. Computation of the Euler characteristic.**

**S2 Code block. Computation of the Betti numbers.**

**S3 Code Block. Computation of Curvature.**

