## Supplemental Code blocks 1-3 for "A hands-on tutorial on network and topological neuroscience"

### S1 Code Block

```python
# Function to compute the Euler Characteristic
def euler(G, verbose=False):
    """Function to compute the Euler characteristic of the clique complex of
    a network

    Parameters
    ----------
    G: networkx graph

    Returns
    -------
    ec: int
      value of the Euler characteristic of the clique complex of the network

    Notes
    -----
    If verbose = True, it will print all the steps of the calculation, so
    that one can check whether the code is working well, or different stages
    of the computation. Otherwise, if verbose=False, only the output will be
    returned.

    """
    def DIAGNOSTIC(*params):
        if verbose:
            print(*params)

    DIAGNOSTIC("Nodes in G: ", G.nodes())
    DIAGNOSTIC("Edges in G: ", G.edges())
    DIAGNOSTIC("Number of nodes: {}, edges: {}".format(G.number_of_nodes(),
               G.number_of_edges()))

    # Compute maximal cliques
    C = nx.find_cliques(G)

    # Create list C with all the cliques
    # Sort each clique, convert it from list to tuple
    C = [tuple(sorted(c)) for c in C]

    DIAGNOSTIC("List with all maximal simplexes/cliques C:",C)
    DIAGNOSTIC("Number of maximal cliques: %i"%(len(C)))


    # Enumerate all simplices/cliques
    S = []
    n = max(len(c) for c in C)

    for k in range(0, n) :

        # Get all (k+1)-cliques, i.e. k-simplices, from max cliques mc
        Sk = sorted(set(c for mc in C for c in itertools.combinations(mc,
                                                                   k+1)))

        DIAGNOSTIC("list of %i-simplex S%i:"%(k,k), Sk)
```

```
        # Check that each simplex is in increasing order
        assert(all((list(s) == sorted(s)) for s in Sk))
        # Assign an ID to each simplex, in lexicographic order
        S.append(dict(zip(Sk, range(0, len(Sk))))) # zip(Sk,range()) is an
        object (composed by tuples) where each element of Sk is associated to
        a number. Then from the zip object create the dictionary where the
        key is the Sk element and the value the number.

    for (k, Sk) in enumerate(S):
        DIAGNOSTIC("Number of {}-simplices: {}".format(k, len(Sk)))
    DIAGNOSTIC("S dictionary", S)

    # The cliques are redundant now
    del C

    # Euler characteristic
    ec = sum(((-1)**k * len(S[k])) for k in range(0, len(S))) # Alternate sum
    of all the simplexes/cliques of different dimensions. len(S[k])
    is how many k-simplex we have. len(S) is the maximum dimension k we can
    find (i.e. the dimension of the simplicial complex)

    DIAGNOSTIC("Euler characteristic:", ec)

    return ec
```

```
# Function to compute the Euler characteristic of the clique complex of the
network with the constraint that we look for cliques up to max dimension k
def euler_k(G, kmax, verbose=False):
    """Function to compute the Euler characteristic of a network with the
    constraint that we look for cliques up to max dimension k

    Parameters
    ----------
    G: networkx graph

    kmax: int
        clique maximum dimension

    Returns
    -------
    Possibilities:
    S: list
     0:Euler characteristic, 1:total cliques, 2:maximum dimension of clique,
     3:number of clique_0, 4:Clique_1, 5:Clique_2, 6:Clique_3, and so on …

    Notes
    -----
    If verbose = True, it will print all the steps of the calculation, so
    that one can check whether the code is working well. Otherwise, if
    verbose=False, only the output will be returned.

    """
    # Prepare maximal cliques
    Nodes = len(G)
    Cliques = nx.find_cliques(G)
```

```
# This function computes the max number for cliques that exists for a
given size
def max_cliques(N, k):
    mclq = 0

    for i in range(0, k):
        mclq += scipy.special.binom(N, k)

    return int(mclq)

Limit = max_cliques(Nodes,kmax)+100 #maximum number of cliques that you
want to find

if verbose == True:
    print("Limit:", Limit)

Cl = []
while True:
    try:
        for i in range(0,Limit):
            clq=next(Cliques)
            if len(clq)<= kmax:
                Cl.append(clq)
    except StopIteration:
        break


# Sort each clique, make sure it is a tuple
C = [tuple(sorted(c)) for c in Cl]
if verbose== True:
    print("C:",Cl)

S = [] # Will contain the number of clique of each order k

for k in range(0, max(len(s) for s in C)):
    # Get all (k+1)-cliques, i.e. k-simplices, from max cliques mc
    Sk = set(c for mc in C for c in itertools.combinations(mc, k+1))
    S.append(len(Sk))

tau = sum(S) # Tau gives the total number of cliques
kmax = len(S) # Kmax is the maximum clique size one can find

if verbose == True:
    print("total # of cliques:", tau)
    print("maximum clique size we can find:", kmax)

Ec = 0 # Ec is the Euler characteristic
Ec = sum(((-1)**i * S[i]) for i in range(0, len(S)))
if verbose== True:
    print("The Euler characteristic EC is:", Ec)

S.insert(0, kmax)
S.insert(0, tau)
S.insert(0, Ec)
```

```
    if verbose== True:
        print("S: Euler, total cliques, maximum dimension of clique, number
              of clique_0,Clique_1,Clique_2, Clique_3")
        print("S:", S)

    # We want to include new elements after kmax with zero, to say that there
    are no simplices with this size - We fixed the elements to 30, this is
    flexible
    for i in range(kmax, 30):
        S.insert(kmax+3, 0)

    return S # The output will be EC, tau, kmax, clique_0,Clique_1,Clique_2,
    Clique_3, and so on…
```

#### S2 Code Block

```
# Function to compute the desired Betti number of the clique complex of the
network
def Betti_k(G, K_input, verbose=False):
    """Function to compute the desired Betti number of a network

    Parameters
    ----------
    G: networkx graph
    K_input: int
           0 if you want to compute Betti-0, 1 if you want to compute Betti-
           1, 2 for Betti-2 etc. Notice that, the higher the K_input, more
           complex/time consuming is the computation.

    Returns
    -------
    B: int
     Betti number

    Notes
    -----
    If verbose = True, it will print all the steps of the calculation, so
    that one can check whether the code is working well. Otherwise, if
    verbose=False, only the output will be returned.

    """
    def DIAGNOSTIC(*params):
        if verbose:
            print(*params)

    DIAGNOSTIC("Nodes in G: ", G.nodes())
    DIAGNOSTIC("Edges in G: ", G.edges())
    DIAGNOSTIC("Number of nodes: {}, edges: {}".format(G.number_of_nodes(),
               G.number_of_edges()))


    # Compute maximal cliques
    C = nx.find_cliques(G)

    # Create list C with all the cliques
    # Sort each clique, convert it from list to tuple
    C = [tuple(sorted(c)) for c in C]

    DIAGNOSTIC("List with all maximal simplex C:", C)
    DIAGNOSTIC("Number of maximal cliques: %i"%(len(C)))

    # Enumerate all simplices

    S = [] # Will be a list of dictionaries

    # Setting the range
    if K_input == 0:
        ini = 0
        fin = 2
    else:
```

```
    ini = K_input-1
    fin = K_input+2

DIAGNOSTIC("I start the loop where I create the required Sk to then
            compute Betti. Sk is a list with the k-simplex")

for k in range(ini, fin): # k has 2 values for betti_0 and 3 values for
betti1_2_3

    Sk = sorted(set(c for mc in C for c in itertools.combinations(mc,
                                                                  k+1)))

    DIAGNOSTIC("list of %i-simplex S%i:"%(k, k), Sk)

    # Check that each simplex is in increasing order
    assert(all((list(s) == sorted(s)) for s in Sk))
    # Assign an ID to each simplex, in order
    S.append(dict(zip(Sk, range(0, len(Sk))))) # zip(Sk,range()) is an
    object (composed by tuples) where each element of Sk is associated to
    a number. Then from the zip object create the dictionary where the
    key is the Sk element and the value the number.


    DIAGNOSTIC("Number of %i-simplices: "%(k), len(Sk))
DIAGNOSTIC("S dictionary", S)

#The cliques are redundant now
del C

# Construct the boundary operator/matrix
# Boundary Matrix
D = [None, None] # List with the two different k-boundary operators

if K_input == 0:
    # D[0] is the zero matrix
    D[0] = (np.zeros((1, G.number_of_nodes()))

for k in range(1, len(S)):

    # Set the index of D[] and the number of nodes in each group for the
    combinatory part
    if K_input == 0:
        index = k
        b = k
    else:
        index = k-1
        b = k+(K_input-1)

    # Create a matrix of  size (len(S[k-1]), len(S[k])
    D[index] = np.zeros( (len(S[k-1]), len(S[k])) )

    for (ks, j) in S[k].items() :

        a = sorted(itertools.combinations(ks, b))

        # Indices of all (k-1)-subsimplices s of the k-simplex ks
        I = [S[k-1][s] for s in sorted(itertools.combinations(ks, b))]
```

```
        for i in range(0,len(I)):
            D[index][I[i]][j] = (-1)**(i)

    if D[index].shape[1] == 0:
        DIAGNOSTIC("I can't create matrix D because the simplicial
                    complex does not have the needed k-simplex")
    DIAGNOSTIC("Boundary matrix:")
    DIAGNOSTIC("D",D[index])
DIAGNOSTIC("D_{} has shape {}".format(K_input, D[0].shape))
DIAGNOSTIC("D_{} has shape {}".format(K_input+1, D[1].shape))

# The simplices are redundant now
del S

# Compute rank and dim(ker) of the boundary operators
# dim(Im)= Rank and dim(ker) = V-rank
rank = [0 if d.shape[1] == 0 else np.linalg.matrix_rank(d) for d in D]
ker = [(d.shape[1] - rank[n]) for (n, d) in enumerate(D)]

#The boundary operators are redundant now
del D
DIAGNOSTIC("ker:", ker)
DIAGNOSTIC("rank:", rank)

# Compute the Betti number
# Betti number
B = ker[0]-rank[1]
DIAGNOSTIC("Betti= ker[0]-rank[1]")
DIAGNOSTIC("End of computation\nBetti %i is:"%K_input, B)

return B
```

### S3 Code Block

```
# Compute nodal curvature based on density
def Curv_density(d, matrix, verbose=False):
    """Compute nodal curvature based on density

    Parameters
    ---------
    d: float
     density value

    matrix: numpy matrix
          connectivity matrix

    Returns
    -------
     curv: numpy array
         array with curvature values

    """

    def DIAGNOSTIC(*params) :
        if verbose : print(*params)
    DIAGNOSTIC("This function run over all nodes and computes the curvature
               of the nodes in the graph")

    # This is the initial Graph
    G = graph_density(d, matrix)  # Filtration function
    temp = Kmaxcliques(G)

    # This lista is a vector V where each v_i is the number of cliques of
size i
    lista = []

    # We suppose that the size of the cliques are smaller than 20, so we
create an empty list of size 20 for the lista
    for i in G.nodes():
        lista.append([0] * 50) # creating a list of lists for each node - all
        empty for the scores for each size for each node

    DIAGNOSTIC("These are all cliques of the Network:")
    DIAGNOSTIC(temp)
    DIAGNOSTIC("We now print the curvature/clique score of each node in the
network")

    # Now we run over all nodes checking if the node belongs to one clique or
another
    # Sc stores the participation rank, which is an earlier step for
computing the curvature
    Sc=[]
    for node in G.nodes(): # now we process for each clique
        score = 0 # This is the initial score of the node in the
        participation rank
```

```
    for clique in temp:
        k = len(clique)
        if node in clique:
            score += 1 # If the node is in the clique raises the score
            lista[node][k-1] += (-1)**(k+1)*1/k # Increases the curvature
            score for a size k with a different weight due to Gauss-
            Bonnet theorem
    Sc.append(score)

    DIAGNOSTIC("The node " + str(node) + " has score =" + str(score))

total = []
for elements in lista:
    total.append(sum(elements)) # This is good if one wants to normalize
    by the maximum
DIAGNOSTIC(total)

# nt is normalized by the sum
# nt2 is normalized by the max
nt=[]
nt2=[]

most = np.argsort(-np.array(total))
for i in most:
        DIAGNOSTIC("The node " + str(i)+ " is in " + str(total[i]) +
                   "cliques")

DIAGNOSTIC("These are the most important nodes ranked according to the
           total clique score")
DIAGNOSTIC(most)
DIAGNOSTIC("These is the array nt")
DIAGNOSTIC(nt)
DIAGNOSTIC("These is the array nt2")
DIAGNOSTIC(nt2)
DIAGNOSTIC("These is the array lista")
DIAGNOSTIC(lista)
DIAGNOSTIC("The output is one vector normalizing the value from the
           maximum")


# curv gives the curvature  - return Sc instead of curv to get the
particiaption rank - notice that you can normalize this quantity in many
ways
curv=[]
for i in range(0, len(lista)):
    curv.append(sum(lista[i]))
curv = np.array(curv)

return curv
```
